# A MICROGLIAL ACTIVITY STATE BIOMARKER PANEL DIFFERENTIATES FTD-GRANULIN AND ALZHEIMER’S DISEASE PATIENTS FROM CONTROLS

**DOI:** 10.1101/2023.06.15.545187

**Authors:** Ida Pesämaa, Stephan A. Müller, Sophie Robinson, Alana Darcher, Dominik Paquet, Henrik Zetterberg, Stefan F. Lichtenthaler, Christian Haass

## Abstract

**Summary:** *Background:* With the emergence of microglia-modulating therapies there is an urgent need for reliable biomarkers to evaluate microglial activation states.

*Methods:* Using mouse models and human induced pluripotent stem cell-derived microglia (hiMGL), which were genetically modified to yield the most opposite homeostatic (*TREM2-* knockout) and disease-associated (*GRN*-knockout) states, we identified microglia activity-dependent markers. Non-targeted mass spectrometry was used to identify changes in microglial and cerebrospinal (CSF) proteome of *Grn*- and *Trem2*-knockout mice. Additionally, we analyzed the proteome of *GRN*- and *TREM2*-knockout hiMGL and their conditioned media. Candidate marker proteins were tested in two independent patient cohorts, the ALLFTD cohort with 11 *GRN* mutation carriers and 12 non-carriers, as well as the proteomic data set available from the European Medical Information Framework Alzheimer’s Disease Multimodal Biomarker Discovery (EMIF-AD MBD).

*Findings:* We identified proteomic changes between the opposite activation states in mouse microglia and cerebrospinal fluid (CSF), as well as in hiMGL cell lysates and conditioned media. For further verification, we analyzed the CSF proteome of heterozygous *GRN* mutation carriers suffering from frontotemporal dementia (FTD). We identified a panel of six proteins (FABP3, MDH1, GDI1, CAPG, CD44, GPNMB) as potential indicators for microglial activation. Moreover, we confirmed three of these proteins (FABP3, GDI1, MDH1) to be significantly elevated in the CSF of AD patients. In AD, these markers differentiated amyloid-positive cases with mild cognitive impairment (MCI) from amyloid-negative individuals.

*Interpretation:* The identified candidate proteins reflect microglia activity and may be relevant for monitoring the microglial response in clinical practice and clinical trials modulating microglial activity and amyloid deposition. Moreover, the finding that three of these markers differentiate amyloid-positive from amyloid-negative MCI cases in the AD cohort suggests that these marker proteins associate with a very early immune response to seeded amyloid. This is consistent with our previous findings in the DIAN (Dominantly Inherited Alzheimer’s Disease Network) cohort, where soluble TREM2 increases as early as 21 years before symptom onset. Moreover, in mouse models for amyloidogenesis, seeding of amyloid is limited by physiologically active microglia further supporting their early protective role. The biological functions of some of our main candidates (FABP3, CD44, GPNMB) also further emphasize that lipid dysmetabolism may be a common feature of neurodegenerative disorders.

*Funding:* This work was supported by the Deutsche Forschungsgemeinschaft (DFG, German Research Foundation) under Germany’s Excellence Strategy within the framework of the Munich Cluster for Systems Neurology (EXC 2145 SyNergy – ID 390857198 to CH, SFL and DP) and a Koselleck Project HA1737/16-1 (to CH).

## Introduction

During the last decade, genome-wide association studies (GWAS) and whole genome sequencing have identified microglial expressed risk variants for late onset Alzheimer’s disease (LOAD).^1^ Triggering receptor expressed on myeloid cells 2 (TREM2) plays a central role in the coordinated switch of microglial activity states upon pathological challenges, and disease risk variants lose their physiologically required activity.^2, 3^ These findings allowed to further elucidate the role of microglia in neurodegenerative disease and initiated the development of microglia modulating therapies.^3^ TREM2 is expressed as a type 1 protein, which is targeted to the cell surface together with its co-receptor DNAX activation protein of 12kDa (DAP12).^4^ On the cell surface TREM2 is proteolytically cleaved to liberate soluble TREM2 (sTREM2). Brain derived sTREM2 is found in biological fluids, such as blood and cerebrospinal fluid (CSF).^5^ Cleavage of TREM2 terminates its cell-autonomous signaling functions.^5^ Since predominantly cell surface TREM2 is cleaved, sTREM2 concentrations in biological fluids are a proxy of signaling-competent full-length TREM2.^6^ sTREM2 levels increase immediately after the initial seeding process when amyloid fibers precipitate and start to form amyloid plaques.^7^ This occurs extremely early during disease progression. In patients with familial AD, sTREM2 increases up to 21 years before disease onset.^7^ Thus, the early TREM2 response of microglia in AD is driven by amyloid precipitation and is apparently a defensive response to a pathological challenge. In fact, seeding of amyloid pathology is boosted in the absence of functional TREM2.^8^ Moreover, increased sTREM2 levels are linked to reduced brain shrinkage and a better cognitive outcome.^9^ Based on these findings, microglia appear to have protective functions at least early during amyloidogenesis. To boost such protective microglial functions, monoclonal antibodies were developed to selectively stabilize and cross-link cell surface TREM2 with the goal to enhance its protective signaling activities.^3, 10^ First clinical trials are already on the way.^3^ TREM2 boosting therapies rely on the development of biomarkers, which allow to monitor target engagement, efficacy of treatment and even stratification of patients with low or high microglial response. Therefore, biomarkers are required, which allow the determination of microglial responses to pathological challenges in human biofluids.

To identify such markers, we made use of models lacking either *TREM2* or the lysosomal protein progranulin (*GRN*) providing us with two completely opposite microglial activation phenotypes. In *TREM2* knock-out (ko) models, microglia are locked in a homeostatic state, while microglia from *GRN* ko models are hyperactivated.^11, 12^ These models therefore capture the two most opposite activity states of microglia and provide optimal models to identify proteins, which reflect either resting or responsive microglial activities. Using mass spectrometry (MS)-based proteomics, we analyzed the proteome of microglia and CSF obtained from *Trem2* ko and *Grn* ko mice as well as cell lysates and media from *TREM2* and *GRN* KO human-induced pluripotent stem cell (iPSC)-derived microglia (hiMGL). Hyperactivated microglia were also reported in patients suffering from frontotemporal dementia (FTD) caused by *GRN* haploinsufficiency (FTD-GRN).^13, 14^ As a first translational approach, we therefore analyzed the CSF proteome of FTD-GRN patients versus healthy non-carriers. This allowed the identification of six proteins as our top biomarker hits, which were consistently increased in hiMGL and in the CSF of *GRN* carriers. We confirmed our potential candidates in AD patients, for which we made use of proteomic data based on human CSF collected as part of the European Medical Information Framework (EMIF)-AD study.^15^ Strikingly, three of the top candidate activity state markers were capable to differentiate amyloid-positive from amyloid-negative individuals diagnosed with mild cognitive impairment (MCI).

## Methods

### Mice

All mice were housed in standard sized individually ventilated cages (IVC), with enriched environment and ad libitum access to food and water. Mice were maintained in a specific pathogen-free facility with a 12-hour light/dark cycle. All experiments and handling of mice was performed in compliance with the German animal welfare law and with approval from the Government of Upper Bavaria. CSF was sampled under the animal license: ROB 55.2-2532.Vet_02-15-69. Afterwards, mice were sacrificed by cervical dislocation followed by brain harvest. CSF and brain tissue were collected from mice of the following mouse strains: C57BL/6J, C57BL/6J *Grn* knockout^16^, and C57BL/6J *Trem2* knockout^17^. The genotyping of these mice was performed as previously described.^13, 16, 17^ Both male and female mice were used for the experiments.

### Mouse CSF sampling

Mice were deeply anesthetized using a mix of medetomidine [0·5 mg/kg], midazolam [5mg/kg], and fentanyl [0·05 mg/kg] (MMF). The MMF-mix was dosed according to bodyweight of each mouse and administrated via intraperitoneal injection. CSF was collected from the cisterna magna by a single stereotactic puncture of the dura using a glass capillary, following a previously described procedure.^18^ The collected CSF samples were centrifuged at 2000 x g for 10 min and visually controlled for blood contamination.

### Microglia isolation from mouse brain – MACS

Immediately after CSF sampling, brains were harvested and microglia were isolated using the Magnetic-Activated Cell Sorting (MACS) technique, following the manufactures instructions (Neural Tissue Dissociation Kit (P), MACS Miltenyi Biotec), with slight modifications similarly to a previously published protocol.^19^ In brief, cerebellum, olfactory bulb, and meninges were removed, and each hemisphere was cut into pieces. Dissected brains were homogenized using an automatic (gentleMACS Dissociator) and enzymatic (Enzyme A (10 µl) + Enzyme P (50 µl)) dissociation process. The homogenates were rinsed through a cell strainer (40 µm, Falcon). Filtered samples were pelleted, washed and resuspended in Hanks’ buffered salt solution (HBSS) supplemented with 7mM HEPES. After washing, cells were magnetically labeled with CD11b (Microglia) Microbeads (Miltenyi Biotec) and diluted in 90 µl MACS buffer (PBS supplemented with 5% BSA). The magnetically labeled cell suspension was loaded onto a MACS LS-column, washed with MACS buffer, and eluted into a Protein LoBind tube. The eluted microglia fraction was pelleted and washed with HBSS to remove any remaining BSA, before all liquid was aspirated and the remaining cell pellet was snap-frozen and stored at -80 °C.

### Human induced pluripotent stem cell (iPSC) culture, CRISPR genome editing and differentiation of iPSC-derived microglia (hiMGL)

All experiments including iPSCs were performed in compliance with all applicable guidelines and regulations. iPSC culture of the female iPSC line A18944 (ThermoFisher, #A18945) based on E8 Flex/VTN (ThermoFisher, #A2858501 and #A14700), CRISPR/Cas9-based genome editing, iPSC quality control measures and differentiation of hiMGL were described previously for the *GRN* ko line^20^ and we followed the same procedures here. With the exception that NEB enzyme SfaNI was used for the analysis with Restriction fragment length polymorphism technique (RFLP) for the *TREM2* ko line, while the MwoI enzyme was used for the *GRN* ko line. We further explain parameters for the generation of *TREM2* ko iPSCs in the Appendix Figure S1.

### Harvesting of hiMGL pellets and conditioned media

Throughout the procedure, cells and PBS were kept on ice. In total, two 6-well plates from each experimental group (WT, *GRN* ko, and *TREM2* ko) were used. The cells were carefully dissociated from the plate using a cell scraper, washed with their media, and subsequently transferred to centrifugation tubes (15 ml), pooled pairwise, yielding n = 6 for each experimental group. The tubes were centrifuged at 3000 x g for 6 min at 4 °C and the conditioned media was transferred to a Protein LoBind tube (Eppendorf) (media fraction). Each well of the plates was washed with 0·5 ml PBS and pooled following the same order as the pooling of the collected cells in the same 15 ml centrifugation tube and the pellet resuspended, resulting in a total volume of 1 ml and n = 6 per experimental group. The 1 ml of PBS with the resuspended cells was transferred to a 1·5 ml Protein LoBind tube (Eppendorf), centrifuged at 3000 x g for 6 min at 4 °C, and PBS aspirated, leaving the pellet. The conditioned media was aliquoted and stored at -80 °C (media fraction). Each cell pellet was stored at -80 °C (microglia fraction).

### Human CSF samples

In total, CSF of 12 symptomatic FTD heterozygous *GRN* mutation carriers (FTD-GRN) and 12 non-symptomatic non-carriers (CON) were analyzed. Data from one *GRN* mutation carrier was excluded as this participant was not fulfilling the criteria of symptomatic FTD, resulting in a comparison of 11 FTD-GRN and 12 CON. All participants were enrolled in ARTFL (U54NS092089) or LEFFTDS (U01AG045390), together ALLFTD, which is a North American research consortium to study sporadic and familiar FTD. ALLFTD has 27 sites in North America. Participants are between the ages of 18-85. Inclusion criteria for ALLFTD are clinical diagnosis of an FTD syndrome, or being a member of a family with a known FTD-associated genetic mutation or a strong family history (modified Goldman score of 1 or 2) of FTD syndromes. For the study presented here, the group defined as FTD-GRN was defined as symptomatic GRN-mutation carriers. All CSF from the ALLFTD consortium were approved and provided by the National Centralized Repository for Alzheimer’s Disease and Related Dementias (NCRAD). CSF obtained from ALLFTD was collected following NCRAD standard operating procedure. In brief, CSF was collected in the morning between 8 am – 10 am, preferably fasted or following a low-fat diet. Lumbar puncture was performed in a sterile field using an atraumatic technique. Within 15 min of collection, CSF samples are centrifuged at 2000 x g for 10 min at room temperature. Post centrifugation CSF is aliquoted into pre-cooled aliquot tubes and within 60 min of CSF collection aliquots are snap frozen on dry ice and stored at -80 °C. Data from the European Medical Information Framework Alzheimer’s Disease Multimodal Biomarker Discovery (EMIF-AD MBD) cohort are publicly available.^15, 21^ Proteomic data from 478 participants were filtered for FABP3, MDH1, GDI1, CAPG, CD44, GPNMB. The abundances of each of the selected proteins were compared across the cohort groups which were defined according to diagnosis as cognitively normal controls (CNC, n = 126), subjective cognitive impairment (SCI, n = 61), mild cognitive impairment (MCI, n = 198), and Alzheimer’s disease (AD, n = 93). The presence or absence of amyloid pathology for the participants of the EMIF-AD MBD cohort was previously assessed by others based on levels of CSF amyloid-β ^21^, which allowed for further stratification of the cohort groups as amyloid-positive (A^+^) and amyloid-negative (A^-^). For correlation analysis CSF levels of TREM2, GFAP and CHI3L1, as included in the publicly available data^15^, were included. The ALLFTD and EMIF-AD MBD are both cross-sectional observational studies.

### Sample preparation for mass spectrometry

In total, 14 sets of samples have been collected from mice, hiMGL and participants of the ALLFTD study. Samples were prepared and analyzed with liquid chromatography tandem mass spectrometry (LC-MS/MS) using a label-free quantification (LFQ) method. The mouse cohort consists out of three mouse lines: Wt, *Grn* ko, and *Trem2* ko – all sacrificed when 12 months old. In total, we analyzed CSF from 14 mice (n(Wt) = 6; n(*Grn* ko) = 4; n(*Trem2* ko) = 4) and microglia from 19 mice (n(Wt) = 7; n(*Grn* ko) = 4; n(*Trem2* ko) = 8). The hiMGL samples included both cell lysates and conditioned media, each collected from three different iPSC lines: WT, *GRN* ko, and *TREM2* ko, with a sample size of n = 6 for each sample type and line. For human CSF, the CSF of 11 symptomatic FTD heterozygous *GRN* mutation carriers (FTD-GRN) and 12 non-symptomatic non-carriers (CON) was analyzed. All samples were prepared and analyzed together according to sample type; mouse CSF, mouse microglia, hiMGL media, hiMGL cell lysates, and hCSF. Mouse CSF was prepared using in-solution digestion. Mouse microglia, and hiMGL media was prepared using filter aided sample preparation (FASP)^22^, while hiMGL cell lysates and hCSF was prepared using Single-pot, solid-phase-enhanced sample preparation (SP3).^23^

### In-solution digestion

From each sample of mouse CSF, a volume of 5 µl was used for in-solution digestion in 0·1% (w/v) sodium deoxycholate according to a previously published protocol.^24^ Peptides were purified by stop and go extraction (STAGE) with C18-packed pipetting tips^25^ and dried using vacuum centrifugation.

### Single-pot, solid-phase-enhanced sample preparation – SP3

15 µl of human CSF was mixed with 15 µl modified STET lysis buffer (150 mM NaCl, 2 mM EDTA pH 8·0, 50 mM Tris pH 7·5, 2% Triton X-100) to inactivate potential viruses. For hiMGL, cell pellets were lysed on ice in STET lysis buffer supplemented with protease inhibitors (Protease Inhibitor Cocktail, Sigma Aldrich, Product P8340). The SP3 sample preparation was performed according to a previously published protocol^26^ using 70% (v/v) acetonitrile for protein binding and four washing steps with 80% (v/v) ethanol. After the sequential digestion with LysC and trypsin (Promega, each protease to protein ratio 1:80), beads were retained with a magnetic rack and samples were filtered with Costar Spin-X spin filters (0·22 µm cut-off) to remove remaining beads, and samples were dried by vacuum centrifugation.

### Sample preparation for mass spectrometry – FASP

Mouse microglia and hiMGL conditioned media were digested using filter aided sample preparation (FASP) as previously described.^22^ For microglia lysates Vivacon spin filters (Sartorius) with 30 kDa cut-off were used, whereas conditioned media samples (500 µl) were concentrated on Vivacon spin filters with 10 kDa cut-off to about 30 µl before starting the protocol. Afterwards, samples were desalted using C18 Stop and Go Extraction (STAGE)^25^ and dried by vacuum centrifugation.

### Liquid chromatography–tandem mass spectrometry – LC-MS/MS

Liquid chromatography-tandem mass spectrometry (LC-MS/MS) was used for label-free quantification (LFQ) of the proteolytic peptides. Two different setups were used. Mouse samples were analyzed on an Easy-nLC 1200 coupled to a Q-Exactive HF mass spectrometer (Thermo Scientific, Waltham, MA, US) as previously described for microglia^19^ using data-independent acquisition and murine CSF samples^27^ applying data dependent acquisition. For human derived samples (hiMGL and hCSF), peptides were separated using a NanoElute nano HPLC equipped with either 30 cm (hCSF samples) or 15 cm (hiMGL samples) 75 µm ID columns packed with ReproSil-Pur 120 C18-AQ, 1·9 µm stationary phase (Dr. Mais GmbH, Germany) applying a 120 min gradient for lysates or a 70 min for hiMGL conditioned media and hCSF, which was online coupled to a TimsTOFpro mass spectrometer (Bruker, Germany). Samples were analyzed using DIA parallel accumulation serial fragmentation. For hCSF, one scan cycle induced one MS1 full scan followed by 2 rows of 50 sequential DIA windows with 18 m/z width for peptide fragment ion spectra with an overlap of 1 m/z covering a scan range of 350 to 1200 m/z (hCSF). The ramp time was fixed to 166 ms and 5 windows were scanned per ramp. This resulted in a total cycle time of 3 s. For hiMGL samples, one scan cycle included one MS1 full scan followed by 2 rows of 40 sequential DIA windows with 22 m/z with for peptide fragment ion spectra with an overlap of 1 m/z covering a scan range of 350 to 1200 m/z. The ramp time was fixed to 100 ms and 4 windows were scanned per ramp. This resulted in a total cycle time of 2·1 s.

### Mass spectrometry data analysis and label free quantification

Database search and label free quantification was performed using the MaxQuant software package (maxquant.org, Max-Planck Institute Munich)^28^ for DDA and Spectronaut (Biognosys, CH) or DIA-NN (https://github.com/vdemichev/DiaNN, Version 1.8).^29^ The analysis of human CSF samples of the ALLFTD study was performed together with CSF samples of other origin than the ALLFTD, for which the data is not presented here. In total, 43 samples were analyzed using the DIA-NN (https://github.com/vdemichev/DiaNN, Version 1.8)^29^, with which the normalization and library generation was performed. As we were specifically interested in FTD and the effect of GRN mutations on the human CSF proteome, we proceeded with the data only obtained from the ALLFTD study. The MS data were searched against canonical fasta databases including one protein per gene of Mus musculus and Homo sapiens from UniProt. For visualizations, unless stated otherwise, proteins are denoted with the gene name. For DIA of mouse microglia, a spectral library generated with samples of microglia with APPPS1 mice^19^ was used for data analysis with Spectronaut. DIA-PASEF data from human CSF and iPSC derived human microglia were analyzed with DIA-NN using a library free search. Trypsin was defined as a protease with cleavage specificity for C-terminal of K and R. Acetylation of protein N-termini and oxidation of methionines were defined as variable modifications. Carbamidomethylation of cysteines was defined as fixed modification. For the database search, two missed cleavages were allowed. False discovery rate (FDR) for both proteins and peptides was adjusted to 1%.

### Statistical analysis

LFQ values were used for relative quantification, log2 transformed and statistically analyzed using Perseus (Version 1.6.15.0)^30^ and GraphPad Prism (Version 9.3.1). Proteins were considered quantifiable when LFQ intensities could be detected in at least three biological replicates per experimental group. No data imputation was performed to replace missing LFQ values. Unless stated otherwise, an unpaired two-tailed Student’s *t*-test was applied to evaluate the significance of proteins with changed abundance. In addition, multiple hypothesis correction was performed using a permutation-based FDR estimation^31^ (threshold: FDR = 0·05, s_0_ = 0·1) or the method of Benjamini, Krieger and Yekutieli^32^ (threshold: FDR = 0·05). Of note, the FDR correction was not considered for the visualization of the data nor the selection of candidates, but is available in the supplementary data. For correlation analysis Spearman correlation was used (two-tail, 95% confidence interval), with the criteria that each selected protein was detected in at least three participants. For multiple comparison (comparing more than two groups), ordinary one-way ANOVA and Tukey’s test were used. Significance was indicated accordingly: “*” = p < 0·05, “**” = p < 0·01, “***” = p < 0·001; “n.s.” = non-significant. Diagnostic performance of each of the panel 6 proteins, as well as for panel 6 as a whole, was assessed using receiver-operating characteristic (ROC) curve analysis and area under the curve (AUC) scores were calculated with 95% confidence using DeLong’s test.

## Results

### Microglial activity state dependent proteomic signatures in mice correlate with transcriptomic signatures and are partially reflected by the CSF proteome

We used a mass spectrometry-based approach to compare the proteome of microglia isolated from 12-months-old *Grn* ko and *Trem2* ko mice and their corresponding age-matched control (Wt) mice (Figure 1A-B). Microglia from 12 months old *Grn* ko mice showed an upregulation of markers associated with microglia activation, such as Apoe, Lgals3, Lyz2, and Clec7a. In contrast, these activation-associated markers were downregulated in the age-matched *Trem2* ko mice (Figure 1A-B). The proteome changes of the isolated microglia matched very well the transcriptomic signatures of the *Grn* ko or *Trem2* ko mice (Figure 1C-D).^11^ This is seen with the microglial activation signature in the *Grn* ko mice, which showed a significant upregulation of Apoe, Clec7a, Cd63, Ctsz, Cd68, Fth1, Ctsb, and Cd9 proteins^11, 12, 33–38^ in both proteomic and transcriptomic datasets (Figure 1E). Similarly, markers indicating a homeostatic microglial signature, such as P2ry12, Cx3cr1, and Sall1 were significantly downregulated on both mRNA and protein level in the *Grn* ko mice (Figure 1F). With the microglial mRNA signature confirmed at the proteome level, we next investigated the proteome of mouse CSF collected pre-mortem from the same mice used for the isolation of microglia. The CSF proteome of the *Grn* ko mice showed several significant changes compared to age-matched Wt. Among the most significantly changed proteins within the CSF proteome of the *Grn* ko, we identified increased levels of Ctsb, Ctsd, Apoe, and Lyz1/2 similar to the microglia proteome (Figure 1G), which is again consistent with their increase at the mRNA level in the DAM signature.^35, 38^ In contrast, the homeostatic signature observed in the *Trem2* ko mouse microglia did not show pronounced changes in the CSF proteome compared to age-matched Wt mice (Figure 1H). In general, the changes observed in the CSF proteome of mice are more subtle than those observed in the proteome of isolated mouse microglia (Figure 1A-B). Nevertheless, DAM-like changes were detected in the CSF and microglial proteome of *Grn* ko mice as indicated by the consistent increase of the DAM markers Apoe, Ctsb, Ctsz, and Lyz2 (Figure 1I).^19, 35, 38^ Noteworthy, besides the DAM-like changes, Chi3l1 (also known as YKL-40), a biomarker for neuroinflammation, showed a significant increase in the CSF of *Grn* ko mice, which is in line with elevated levels observed in the CSF of humans with neurodegenerative disorders (Figure 1G).^39, 40^

**Figure 1:**
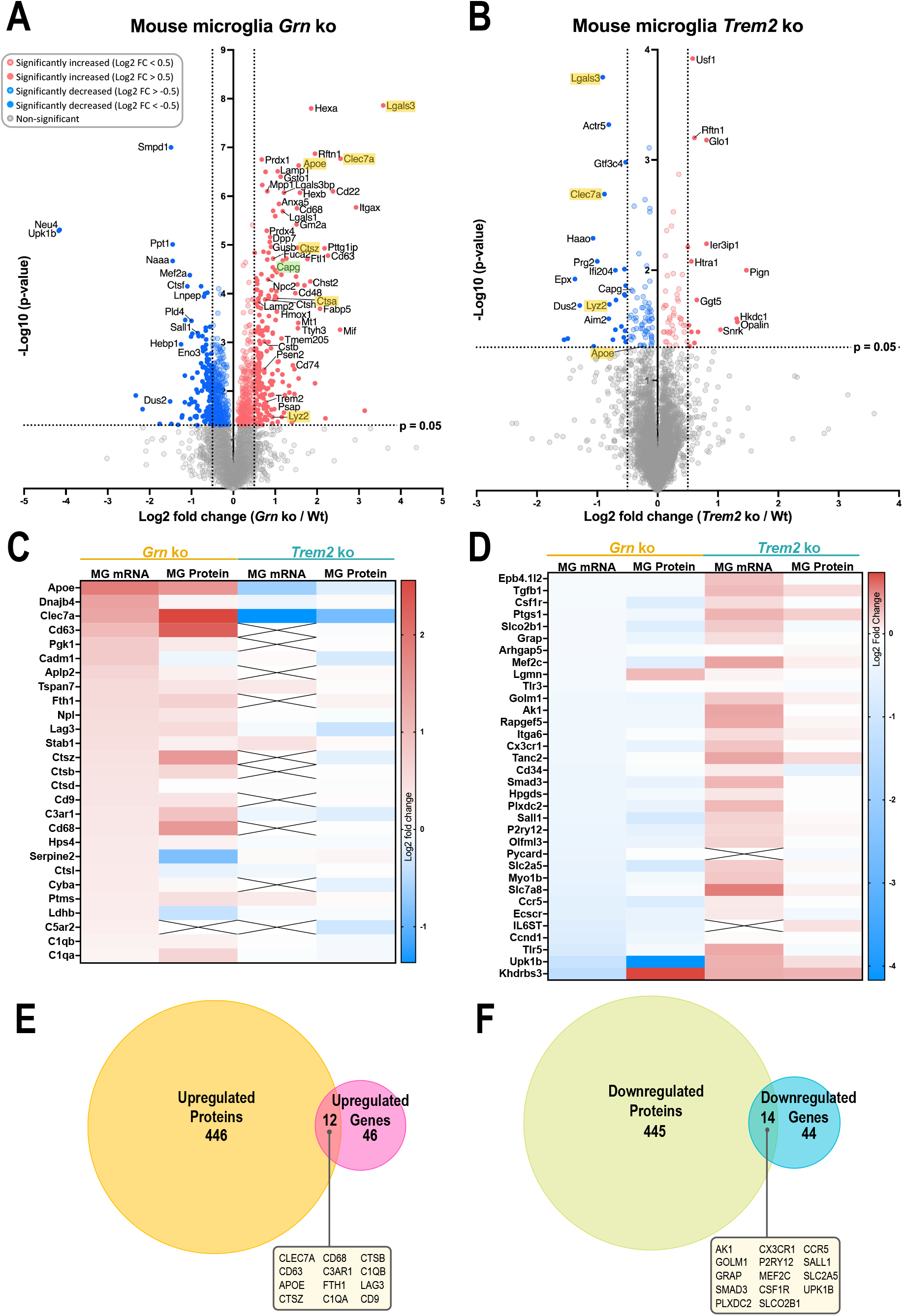
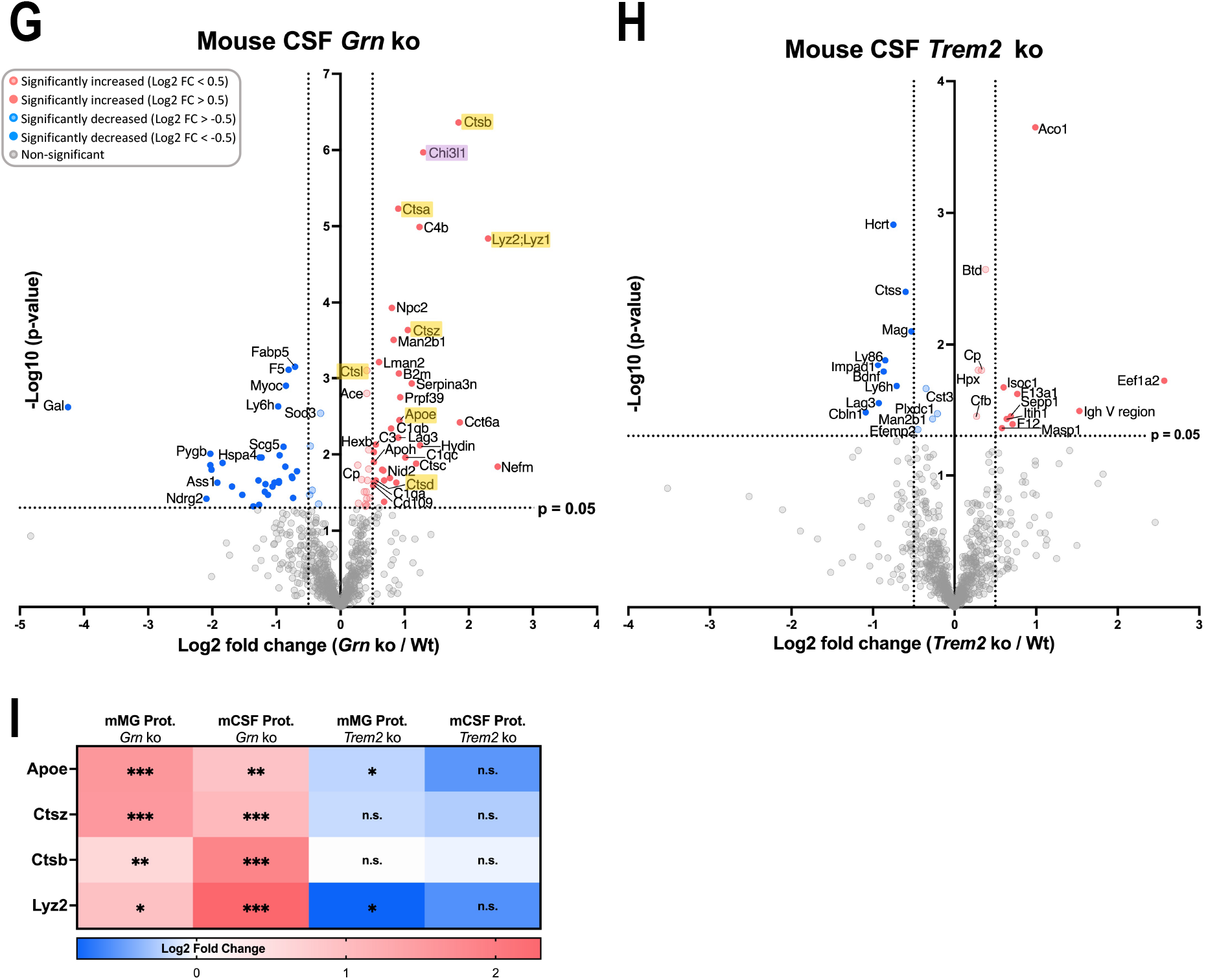
Microglial activation markers in *Grn* ko and *Trem2* ko are consistently changed on mRNA and protein level in mice and partially reflected in the mouse CSF proteome. **(A-B)** Volcano plots showing upregulated (red) and downregulated (blue) proteins in microglia isolated from mice with the following genotypic comparisons: **(A)** *Grn* ko / Wt and **(B)** *Trem2* ko / Wt. Selected cut-off values: p-value < 0·05 (colored dots) and -0·5 > Log2 FC > 0·50 (filled dots). The -log10 transformed p-value of each protein is plotted against its log2 fold change. **(C-D)** Log2 fold-changes (relative to age-matched Wt) of microglial expressed mRNAs and related proteins, show very similar transcriptomic and proteomic changes. mRNA signatures were generated in independent samples from Cd11b-positive microglia isolated from 5·5 months old *Grn* ko and *Trem2* ko mice, respectively.^11^ Crosses indicate missing values. **(E-F)** Comparing microglial proteome and transcriptome of *Grn* ko mice. **(E)** Significantly upregulated proteins (orange) versus significantly upregulated genes (pink). Overlap of 12 markers significantly upregulated on both protein and mRNA level (text box). **(F)** Significantly downregulated proteins (green) versus significantly downregulated genes (blue). Overlap of 14 markers significantly downregulated on both protein and mRNA level (text box). Selected cut-off values: p-value < 0·05. **(G-H)** Comparing upregulated (red) and downregulated (blue) proteins in mouse CSF with the following genotypic comparisons: **(G)** *Grn* ko / Wt and **(H)** *Trem2* ko / Wt. Selected cut-off values: p-value < 0·05 and -0·5 > Log2 FC > 0·50. **(I)** Changes (relative to age-matched Wt) of Apoe, Ctsz, Ctsb, and Lyz2 are consistent in the proteome of isolated microglia and in the proteome of CSF sampled from the same mice. Statistical differences were calculated using Student’s t-test. * = p-value < 0·05, ** = p-value < 0·01, *** = p-value < 0·001; n.s. = non-significant. False discovery rate (FDR) was not considered for the presented visualizations. All presented proteomic data was obtained by LC-MS.

### Proteomic signatures of *Grn* ko mouse microglia correlate with human iPSC derived *GRN* ko microglia

To examine the potential translational applicability of our proteomic results from mouse microglia and CSF, we investigated proteomic changes in cell lysates and conditioned media of hiMGL harboring a *GRN* ko or *TREM2* ko (Appendix Figure S2).^20^ In line with the results obtained with mouse microglia, a larger number of proteins were significantly upregulated in the *GRN* ko hiMGL compared to *TREM2* ko hiMGL (Figure 2A-B). Cathepsins (CTS) including CTSA and CTSZ, but also APOE and LGALS3 were significantly upregulated in the *GRN* ko hiMGL compared to WT control, which is in line with our findings in the microglia proteome of *Grn* ko mouse (Figure 1A and 2A).

**Figure 2:**
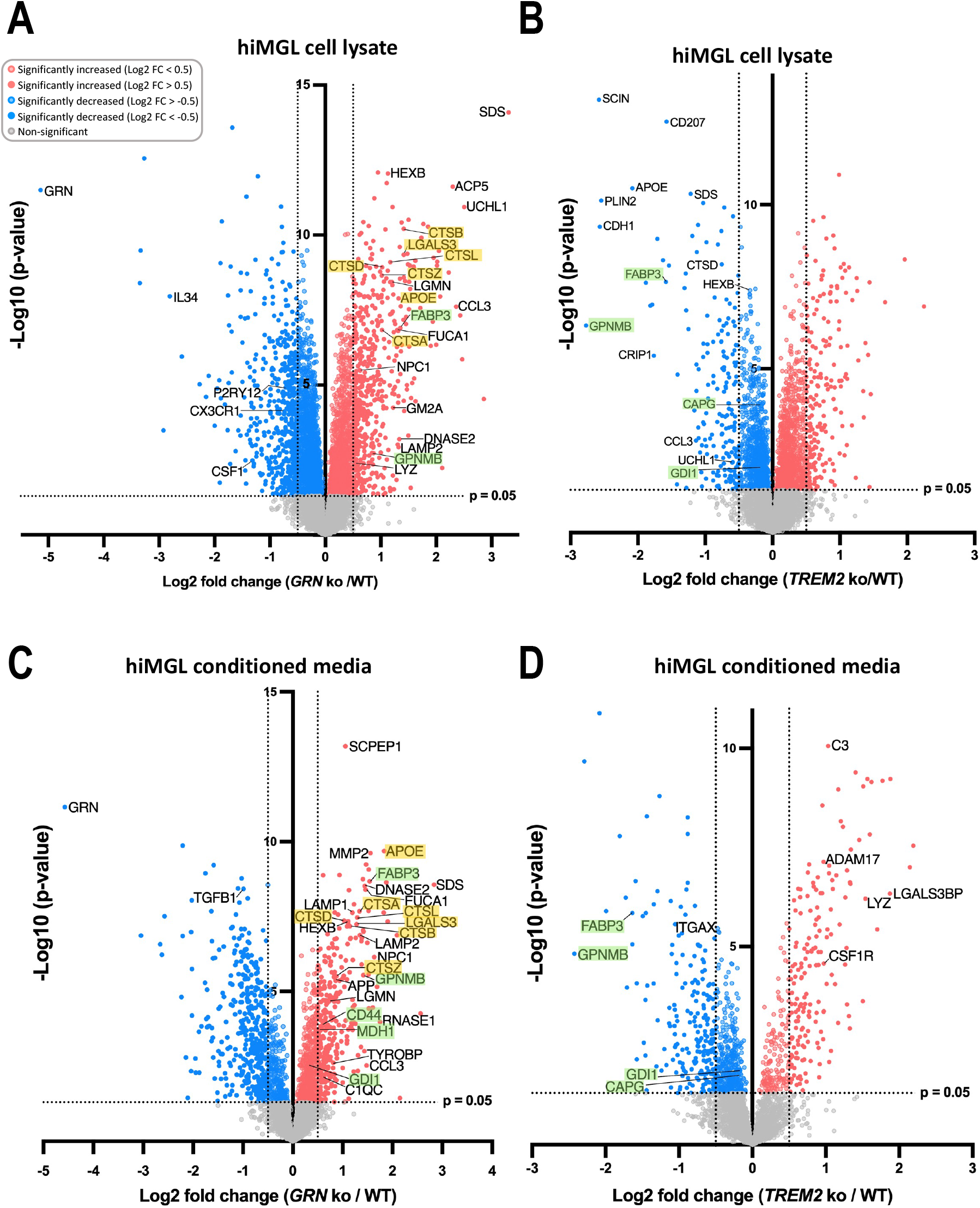
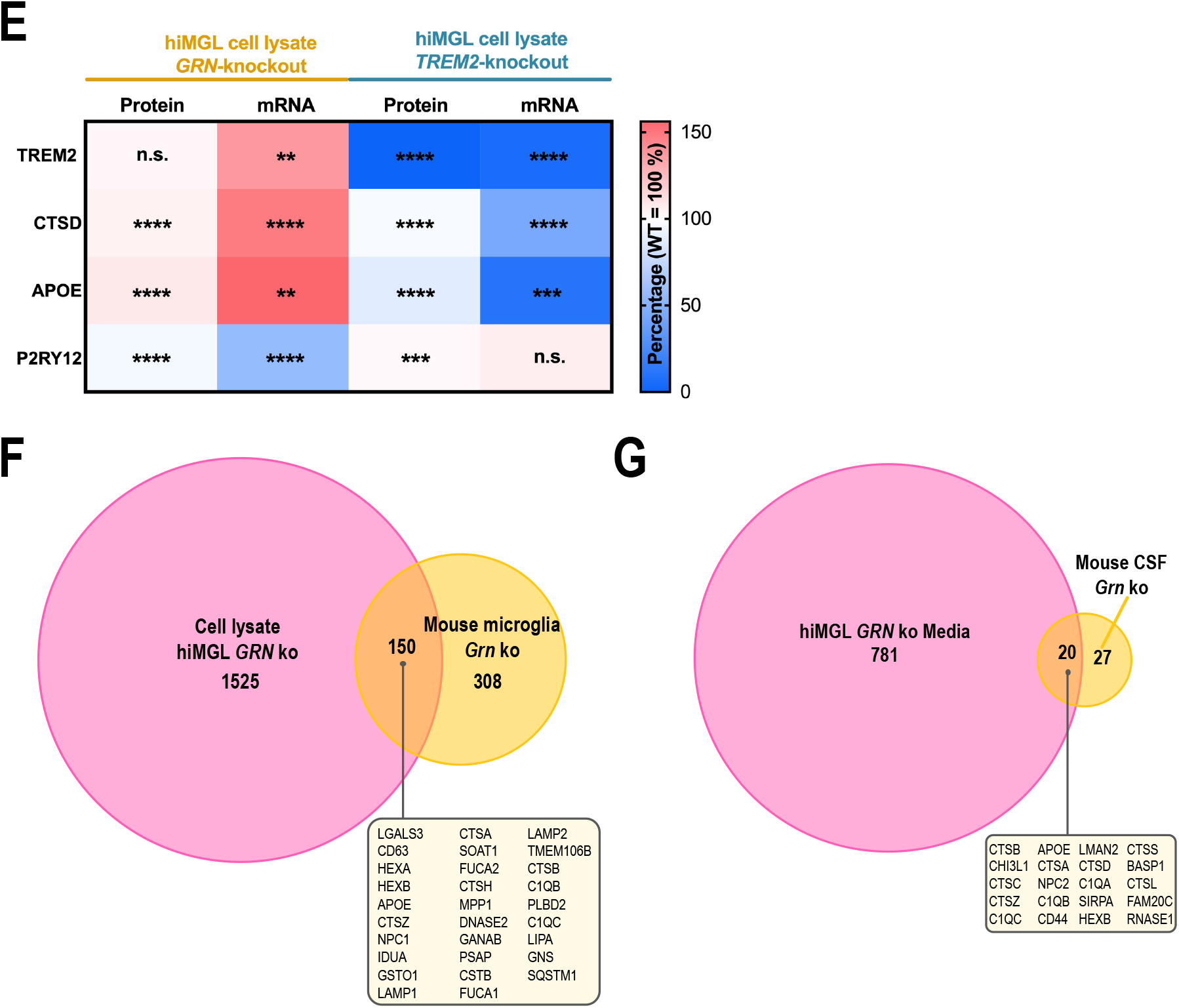
The protein signature of activated mouse microglia is comparable with hiMGL and associates well between mouse CSF and the hiMGL secretome. **(A-B)** Comparing upregulated proteins (red) versus downregulated proteins (blue) with significant changes indicated as colored dots, in the proteome of iPSC-derived microglia (hiMGL) with the following genotypic comparisons: **(A)** *GRN* ko / WT and **(B)** *TREM2* ko / WT. (**C-D)** Comparing proteomic changes in conditioned media from hiMGL (secretome) with the following genotypic comparisons: **(C)** *GRN* ko / WT **(D)** *TREM2* ko / WT. Cathepsins, as well as APOE and LGALS3 are marked in yellow. Other proteins of interest are marked in green. Selected cut-off values: p-value < 0·05 (colored dots) and -0·5 > Log2 FC > 0·50 (filled dots). The -log10 transformed p-value of each protein is plotted against its log2 fold change. **(E)** Marker proteins related to microglia activation showed significant consistency between the proteome (first and third column) and transcriptome (second and fourth column) in hiMGL cell lysates. mRNA data were obtained by qPCR. **(F)** Proteins significantly (p-value < 0·05) upregulated in lysates of hiMGL *GRN* ko (pink) and *Grn* ko mice (orange). 150 proteins were significantly upregulated in both models (a selection of overlapping proteins is shown in the box). **(G)** Proteins significantly (p-value < 0·05) upregulated in the conditioned media of *GRN* ko hiMGL (pink) and CSF sampled from 12 months old *Grn* ko mice (orange). 20 proteins were significantly upregulated in both mouse CSF and conditioned media of hiMGL (box). Statistical differences were calculated using Student’s t-test. * = p-value < 0·05, ** = p-value < 0·01, *** = p-value < 0·001; n.s. = non-significant. False discovery rate (FDR) was not considered for the presented visualizations. Mice used for analysis of the proteome were 12 months old. All presented proteomic data was obtained by LC-MS.

Next, we analyzed the hiMGL secretome (conditioned media) (Figure 2C-D). The proteomic signature observed in lysates of *GRN* ko hiMGL was reflected in conditioned media of *GRN* ko hiMGL. For example, several cathepsins (CTSA, CTSB, CTSD, CTSL, and CTSZ) were significantly increased in lysates and media of *GRN* ko hiMGL (Figure 2A and C). Furthermore, the hiMGL models showed a robust correlation between the microglial transcriptome and proteome. For example, proteins with strong association to microglial activation, such as TREM2, APOE and CTSD were all upregulated on both, mRNA and protein level in the *GRN* ko hiMGL, compared to WT (Figure 2E and Appendix Figure S2A). Consistent with the activation state of *GRN* ko hiMGL, levels of the homeostatic marker P2RY12 were significantly reduced in the *GRN* ko hiMGL transcriptome as well as the proteome (Figure 2E and Appendix Figure S2A). In contrast, P2RY12 protein levels are slightly increased in the *TREM2* ko hiMGL, which are locked in a homeostatic state (Figure 2E and Appendix Figure S2A).^11, 12, 20^ The microglial proteome signatures observed in mice were comparable with the changes measured in hiMGL. In total, 33% of proteins significantly upregulated in *Grn* ko mouse microglia were also significantly upregulated in the cell lysate of *GRN* ko hiMGL (Figure 2F). In the secretome, > 42% of proteins significantly upregulated in CSF of *Grn* ko mice were also significantly upregulated in the secretome of *GRN* ko hiMGL (Figure 2G). This suggests that the proteomic signature for microglial activation that we capture with our mouse models are at least in part conserved between mouse and human. As in our mouse models, the changes in the *TREM2* ko hiMGL model are more subtle than those in *GRN* ko hiMGL. Comparison of the number of significantly upregulated proteins in cell lysates of *GRN* ko hiMGL (1675 proteins (Figure 2F)) versus significantly upregulated proteins in lysates of *TREM2* ko hiMGL (1247 proteins (Appendix Figure S2B), reveals that the *GRN* ko hiMGL have 34% more proteins with significantly increased abundance compared to the *TREM2* ko hiMGL. With a lower number of significantly upregulated proteins in the *TREM2* ko models, the overlaps are also limited when comparing the microglia proteome of *Trem2* ko mice with *TREM2* ko hiMGL, resulting in an overlap of only 5 proteins (GLO1, PRPS2, HPRT1, SORT1, and DHX36) (Appendix Figure S2B). Of note, no proteins overlapped when comparing the 13 significantly increased proteins in the *Trem2* ko mouse CSF versus the 316 proteins with higher abundance in the secretome of *TREM2* ko hiMGL (Appendix Figure S2C).

### Consistent changes in the CSF proteome of symptomatic *GRN* mutation carriers and *GRN* ko hiMGL

For further validation of our proteomic results from mice and hiMGL, we analyzed the CSF proteome of human *GRN* mutation carriers. In total, the cohort included 11 symptomatic heterozygous *GRN* mutation carriers (FTD-GRN; MC = mutation carrier), of which 3 patients were diagnosed with mild cognitive impairment (MCI) at the timepoint of CSF sampling. In addition, the CSF of 12 non-symptomatic controls (CON; NC = non-carrier) without *GRN* mutations were included (Appendix Table S1). As expected, GRN was significantly less abundant in the CSF of FTD-GRN patients compared to healthy controls (Figure 3A). On the other hand, levels of neurofilament light (NEFL) were significantly increased in the CSF of FTD-GRN patients, indicating massive neurodegeneration (Figure 3A). Comparison of the CSF proteome of FTD-GRN patients to the proteome data obtained from conditioned media of *GRN* ko hiMGL secretome provided the identification of 26 proteins with increased abundance in both fluids (p-value < 0·05, not considering FDR-correction) (Figure 3B-C). When comparing these 26 proteins with our proteomic data derived from the *Grn* ko mouse microglia, a statistically significant upregulation is observed for four proteins, which therefore show a consistent increase across species and models: FUCA1, HEXB, FUCA2, and CAPG. In the CSF of *Grn* ko mice, the increased levels of three proteins found in the human CSF also reached a statistical significance: HEXB, CD44, and CHI3L1 (Appendix Figure S3).

**Figure 3:**
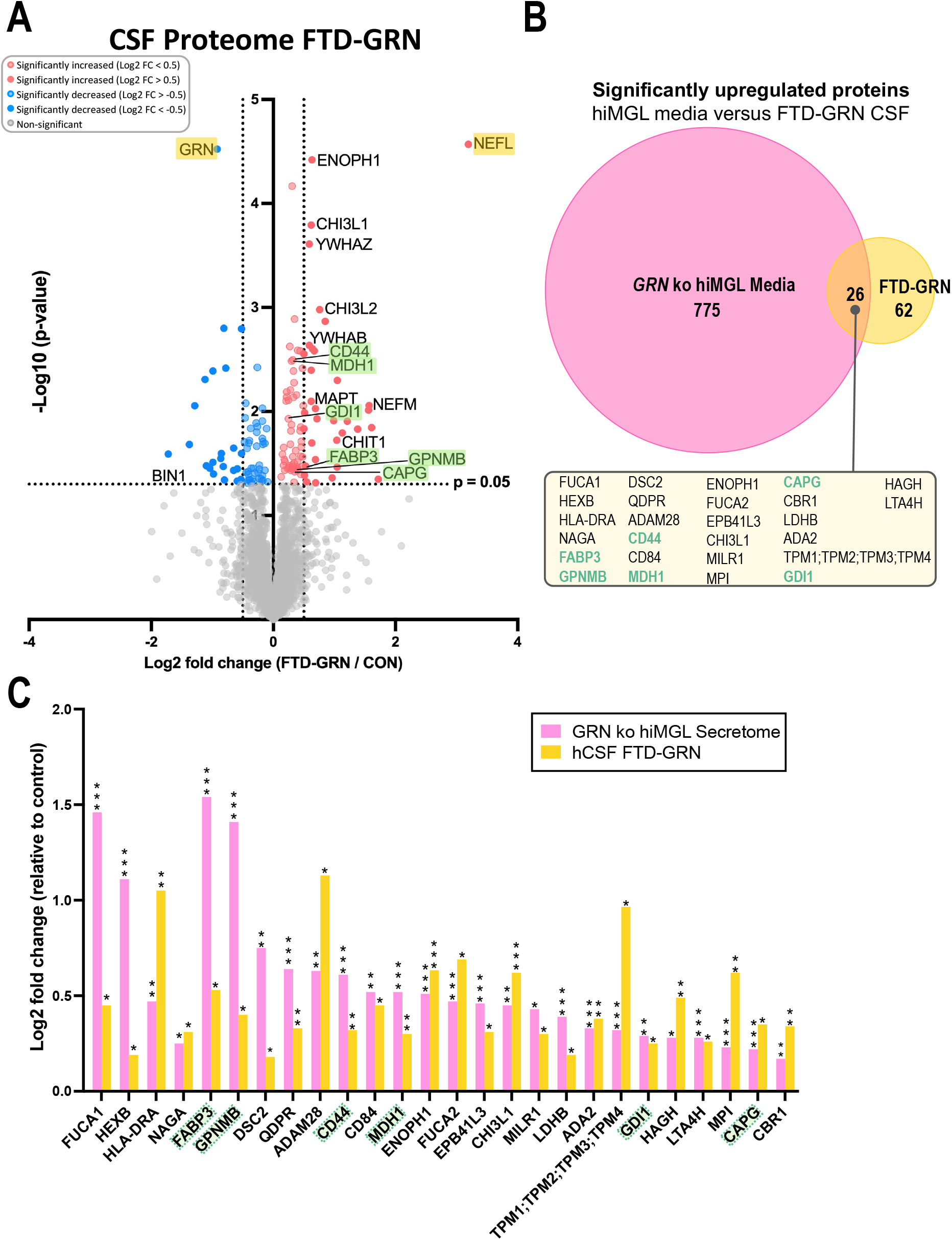
Overlapping proteomic changes observed in the CSF proteome of symptomatic GRN carriers and the secretome of GRN ko hiMGL. (**A)** Comparison of the CSF proteome of symptomatic heterozygous *GRN* mutation-carriers (FTD-GRN) and healthy controls (CON). Significantly upregulated proteins (red) versus significantly downregulated proteins (blue) are shown. Selected cut-off values: p-value < 0·05 (colored dots) and -0·5 > Log2 FC > 0·50 (filled dots). The -log10 transformed p-value of each protein is plotted against its log2 fold change. (**B)** Identification of microglial activity state-dependent proteins by comparing the significantly upregulated proteins in human CSF of FTD-GRN patients (orange) to the significantly upregulated proteins detected in the conditioned media of *GRN* ko hiMGL (pink). In the CSF of FTD-GRN patients, 88 proteins were significantly upregulated compared to healthy non-carriers. 26 (30%) of these proteins were also detected as significantly upregulated in the conditioned media of hiMGL lacking *GRN* compared to media of WT hiMGL (box). **(C)** Quantitative comparison of the 26 proteins detected as significantly upregulated in both, the human CSF (hCSF) of FTD-GRN patients (orange) and the media of *GRN* ko hiMGL (pink) presented as Log2 Fold Change in relation to respective control (CON and WT, respectively). Proteins of interest are marked in yellow and green. Statistical differences were calculated using Student’s t-test. * = p-value < 0·05, ** = p-value < 0·01, *** = p-value < 0·001, n.s. = non-significant. False discovery rate (FDR) was not considered for the presented visualizations.

### Identification of six microglia activation-dependent proteins

By comparing the 26 proteins that were upregulated in both the secretome of *GRN* ko hiMGL and in the CSF of FTD-GRN patients (Figure 3B and Appendix Table S2) we identified six proteins, which we refer to as panel 6, including fatty acid binding protein 3 (FABP3), malate dehydrogenase 1 (MDH1), GDP dissociation inhibitor-1 (GDI1), macrophage-capping protein (CAPG), CD44, and glycoprotein NMB (GPNMB) (Figure 3A-C and 4A). All proteins of panel 6 were significantly increased in the CSF of FTD-GRN patients and in the conditioned media of *GRN* ko hiMGL. FABP3 and GPNMB are the only proteins from panel 6 that were also significantly upregulated in the cell lysate of the *GRN* ko hiMGL. In *Grn* ko mice, CD44 was the only protein within panel 6 that was significantly upregulated in CSF, while CAPG was the only protein significantly upregulated in mouse microglia (Figure 4B-C and Appendix Figure S3). Investigating each of the six proteins separately, within the *GRN* ko hiMGL secretome and the CSF of FTD-GRN patients, FABP3 stands out with increasing CSF levels of > 50% in FTD-GRN patients compared to healthy controls (Appendix Figure S4A). Noteworthy, the FABP3, MDH1 and GDI1 CSF levels in the three *GRN* mutation carriers with less prominent symptoms and diagnosed with MCI, are below average of the FTD-GRN group (Appendix Figure S4A-C). CSF levels of the remaining three proteins (CAPG, CD44, and GPNMB) were significantly elevated in *GRN* mutation carriers, but without any reducing effect caused by the MCI diagnosis (Appendix Figure S4D-F).

**Figure 4:**
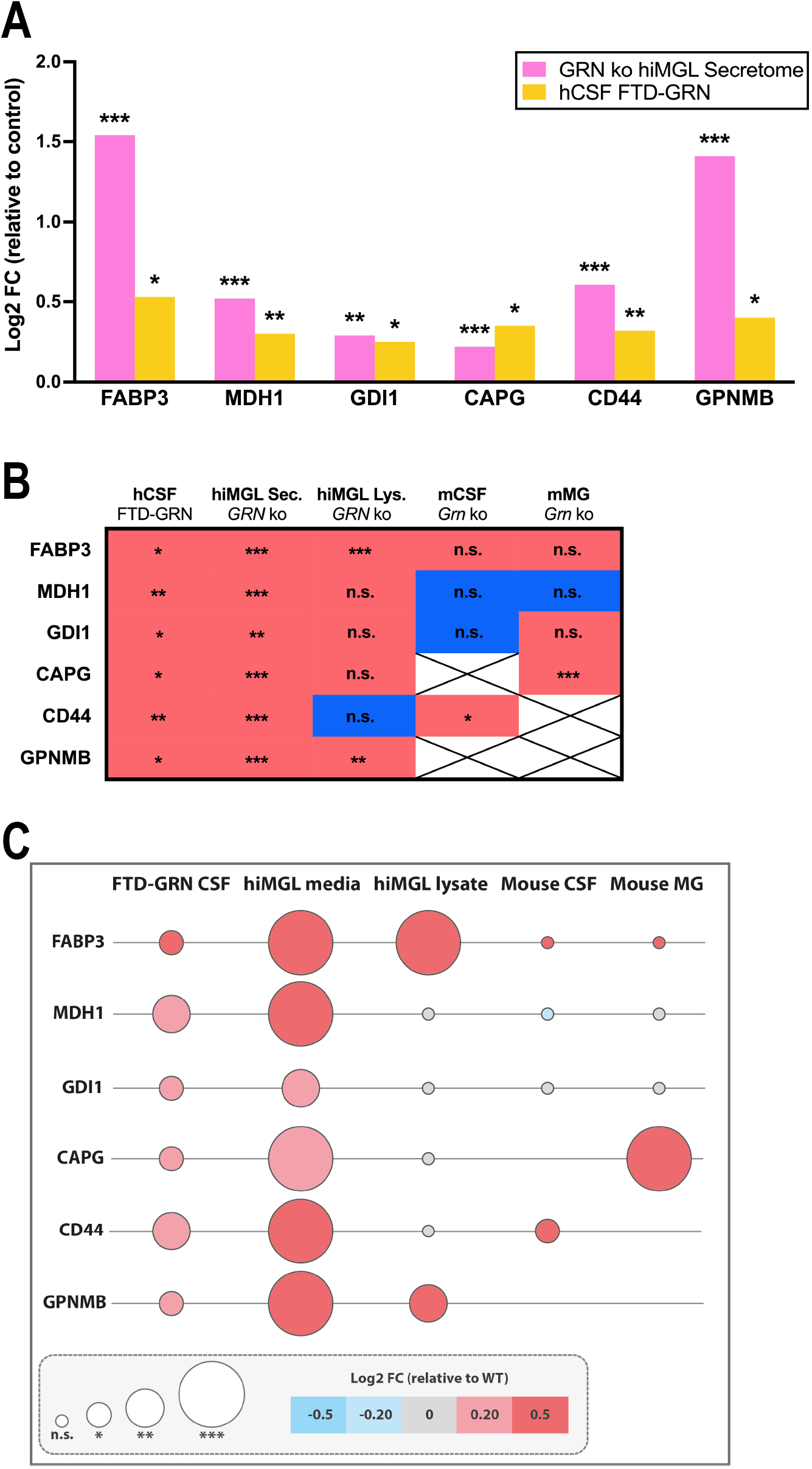
Identification of six microglial activation-dependent proteins. (**A)** Log2 transformed abundances of panel 6 proteins (FABP3, MDH1, GDI1, CAPG, CD44, and GPNMB) in the secretome of *GRN* ko hiMGL (pink) and in CSF of symptomatic FTD-GRN patients (orange). The abundance of each group is normalized to the abundances measured in control groups, secretome of WT hiMGL and CSF levels of healthy non-carrier (CON). **(B)** Binary representation of the up-(red) and downregulation (blue) of each protein in hCSF of FTD-GRN patients, conditioned media of *GRN* ko hiMGL, cell lysates of *GRN* ko hiMGL, CSF (mCSF) of *Grn* ko mice, and microglia (mMG) of *Grn* ko mice. Crosses indicate missing values. **(C)** Comparison of Log2 transformed abundance of panel 6 proteins in CSF from FTD-GRN patients, conditioned media from *GRN* ko hiMGL, *GRN* ko hiMGL cell lysates, CSF and microglia from *Grn* ko mice. Circle size indicates statistical significance based on p-value obtained from Student’s t-test, * = p-value < 0·05, ** = p-value < 0·01, *** = p-value < 0·001, n.s. = non-significant. False discovery rate (FDR) was not considered for the presented visualizations. Color intensity indicates the degree of up-(red) or downregulation (blue). Unchanged abundances are indicated in gray and are defined by -0·2 < Log2 fold change < 0·2.

### Panel 6 proteins distinguish FTD-GRN patients from controls and correlate with levels of CHI3L1

The diagnostic power of panel 6, to distinguish FTD-GRN patients from healthy controls was evaluated using ROC curve analysis. When combined, the panel 6 proteins generated an AUC of 0·87 (95% CI = 0·64 to 1·0) (Figure 5A). Notably, our results revealed that FABP3 alone generated an AUC of 0·93 (95% CI = 0·78 to 1·0) (Figure 5A-B).

**Figure 5:**
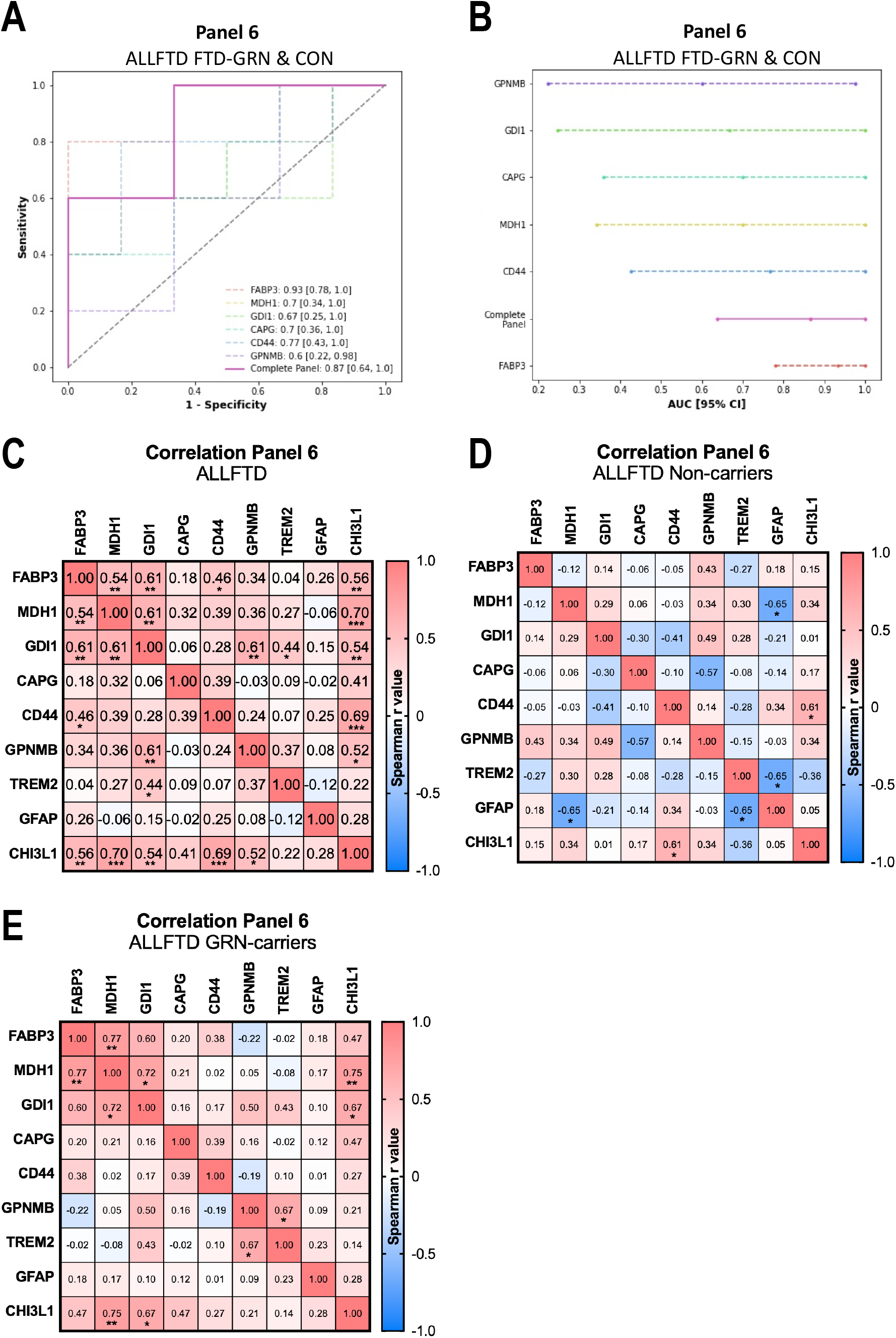
Panel 6 proteins successfully distinguishing FTD-GRN patients from controls and correlate with levels of CHI3L. **(A-B)** ROC curve analysis indicating the diagnostic performance of panel 6 (AUC = 0·87, 95% CI = 0·64 to 1·0) to separate cohort participants as affected (FTD-GRN) versus not affected (CON). **(C-E)** Correlation analysis of CSF levels of panel 6 proteins and TREM2 (microgliosis), GFAP (astrogliosis), and CHI3L1 (also known as YKL-40, neuroinflammation) within **(C)** the entire ALLFTD cohort (n (ALLFTD) = 23)), within **(D)** the control group only (n (NC) = 12), and within **(E)** the symptomatic FTD-GRN group (n (MC) = 11). CSF levels of TREM2 (microgliosis), GFAP (astrogliosis), and CHI3L1 (also known as YKL-40, neuroinflammation) are included based on the quantitative data obtained from LC-MS analysis. Spearman r values, correlations are categorized as strong correlation (Spearman r > 0·5) or very strong (Spearman r > 0·75). * = p-value < 0·05, ** = p-value < 0·01, *** = p-value < 0·001, n.s. = non-significant. FDR was not considered for the presented visualizations.

For further characterization of panel 6, the correlation between each of the six proteins was investigated within the entire cohort (Figure 5C) as well as within each group (healthy non-carriers and FTD-GRN) (Figure 5D-E). Although not significantly altered in the CSF data, we included known markers for microgliosis (TREM2), astrogliosis (GFAP), and neuroinflammation (CHI3L1, also known as YKL-40) for correlation analysis. A significant correlation was observed between FABP3, MDH1, and GDI1 (Figure 5C). The levels of these proteins also significantly correlated with the levels of CHI3L1 (Figure 5C). In general, the observed correlations were weaker in the non-carriers compared to the *GRN* mutation carriers (Figure 5D-E). Strong correlations (defined by Spearman r > 0·5) were observed between GPNMB-CHI3L1, GDI1-GPNMB, GDI1-CHI3L1, FABP3-MDH1, FABP3-CHI3L1, MDH1-CHI3L1, CD44-CHI3L1, GDI1-MDH1, and FABP3-GDI1 (Figure 5C). Very strong correlations (defined by Spearman r > 0·75) were observed between FABP3-MDH1 and MDH1-CHI3L1 in the FTD-GRN group (Figure 5E).

### CSF levels of FABP3, MDH1, and GDI1 are significantly elevated in AD and mild cognitive impairment and driven by Aβ

For further validation, we investigated the abundances of the six selected proteins in a previously published proteomic dataset from the CSF of 478 participants of the EMIF-AD MBD cohort.^15^ Within this cohort three of our six selected proteins were significantly altered between cohort groups stratified according to diagnosis, namely FABP3, MDH1, and GDI1 (Figure 6A-B). The CSF levels of CAPG, CD44, and GPNMB did not differ between the cohort groups (Appendix Figure S5A-B). Of note, the CSF levels of FABP3, MDH1, and GDI1 were not only significantly elevated in participants diagnosed with AD, but also in the non-dementia classed MCI group when compared to CNC (Figure 6B). Stratification according to amyloid status allowed further separation within the MCI and AD group. CSF levels of FABP3, MDH1, and GDI1 were significantly different within the MCI group, with a highly significant elevation in MCI cases with amyloid abnormalities compared to amyloid-negative MCI cases (Figure 6C-D). Furthermore, CSF levels of MDH1 became significantly different within the AD group, when stratifying this group according to amyloid status (Figure 6D).

**Figure 6:**
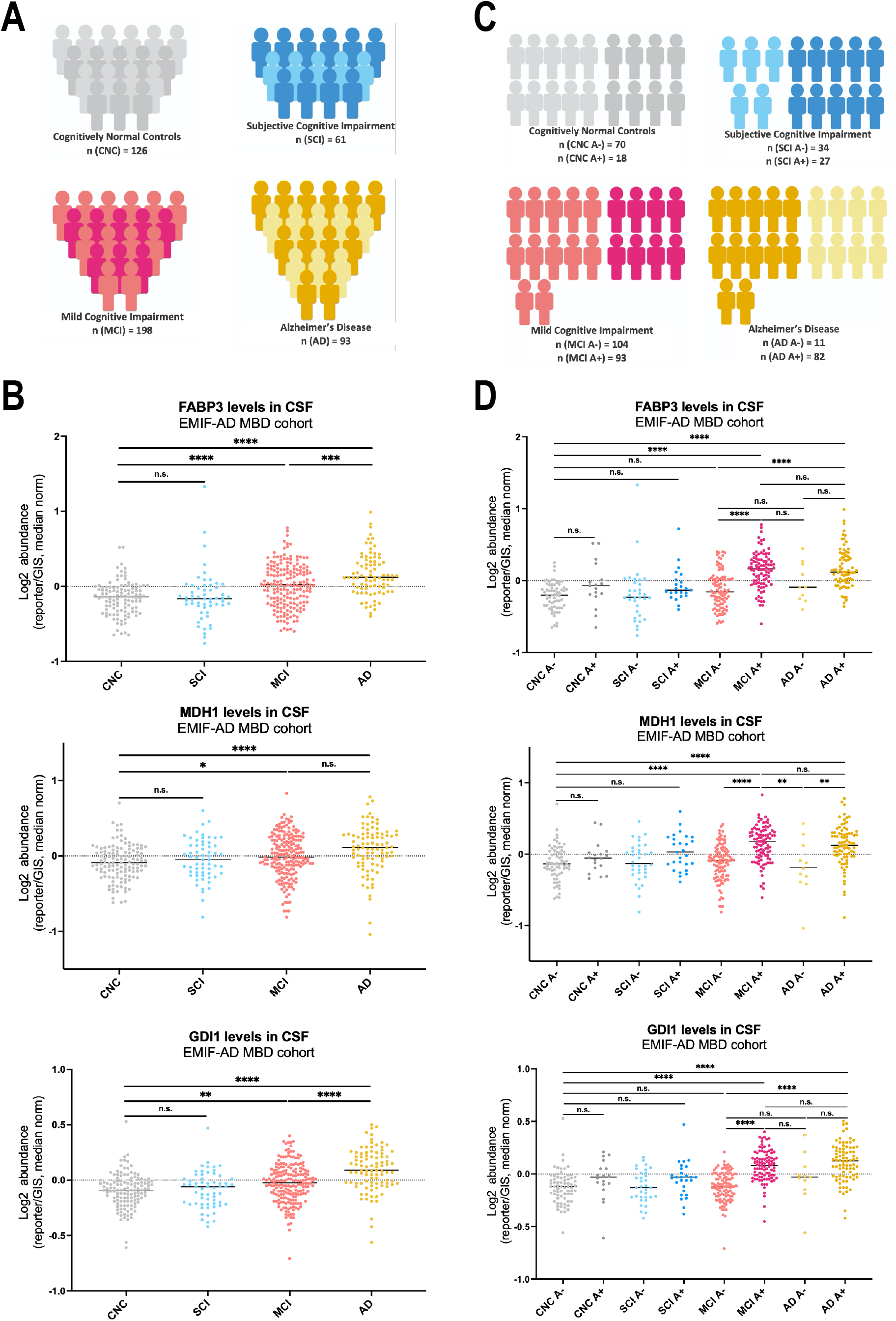
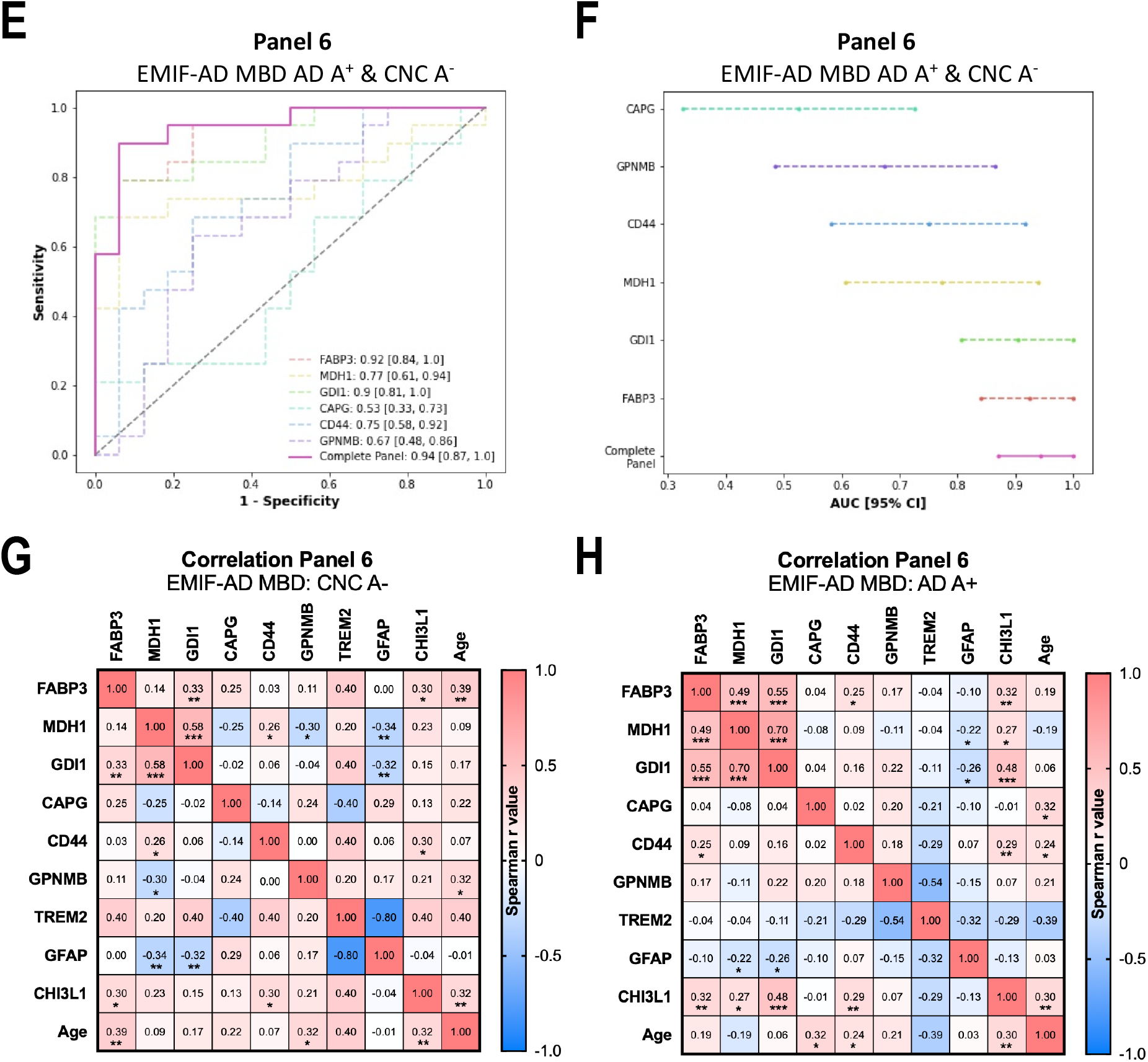
CSF levels of FABP3, MDH1, and GDI1 correlates with each other and are significantly elevated in an amyloid dependent manner in MCI and AD. (**A)** Overview of the EMIF-AD MBD cohort with stratification according to diagnosis only. **(B)** CSF levels of FABP3 (top), MDH1 (middle), and GDI1 (bottom) in cognitively normal controls (CNC) (gray), SCI (blue), MCI (pink), and AD (yellow). CSF levels are defined by Log2 abundance, normalized to the abundance measured in the TMT-reference control channel^15^ and the median of each individual. **(C)** Overview of the EMIF-AD MBD cohort with stratification according to diagnosis and amyloid status as assessed by the levels of CSF amyloid-β_1-42_ ^21^. **(D)** CSF levels of FABP3 (top), MDH1 (middle), and GDI1 (bottom) in CNC A^-^ (light gray), CNC A^+^ (dark gray), SCI A^-^ (light blue), SCI A^+^ (dark blue), MCI A^-^ (light pink), MCI A^+^ (dark pink), AD A^-^ (light yellow), and AD A^+^ (dark yellow). A^-^ = amyloid normal, A^+^ = amyloid abnormal.^15^ Ordinary one-way ANOVA (alpha = 0·05). **(E-F)** ROC curve analysis indicating the diagnostic performance of panel 6 (AUC = 0·94, 95% CI = 0·87 to 1·0) to separate cohort participants as affected (amyloid-positive AD) versus not affected (amyloid-negative CNC). (**G-H)** Correlation analysis of panel 6 proteins and TREM2 (microgliosis), GFAP (astrogliosis), and CHI3L1 (also known as YKL-40, neuroinflammation) in **(G)** amyloid-negative CNC and in **(H)** amyloid-positive AD patients. CSF levels are defined by Log2 abundance, normalized to the abundance measured in the TMT-reference control channel and the median of each individual. Each cell is colored and labeled according to Spearman r value. * = p-value < 0·05, ** = p-value < 0·01, *** = p-value < 0·001, n.s. = non-significant. FDR was not considered for the presented visualizations.

The diagnostic power of panel 6, to distinguish amyloid-positive AD patients from amyloid-negative healthy controls was evaluated using ROC curve analysis. When combined, the panel 6 proteins generated an impressive AUC value of 0·94 (95% CI = 0·87 to 1·0) (Figure 6E). Interestingly, our results revealed that FABP3, GDI1, and MDH1 are the best performing proteins among the six panel proteins, with AUC values of 0·92 (95% CI = 0·84 to 1·0), 0·90 (95% CI = 0·81 to 1·0), and 0·77 (95% CI = 0·61 to 0·94), respectively (Figure 6E-F). Finally, we found a similar correlation between FABP3, MDH1 and GDI1, as observed in the ALLFTD cohort (Figure 5C and Figure 6 G-H). The observed correlation was stronger in the group of amyloid-positive AD patients compared to the amyloid-negative control group (CNC) (Figure 6G-H). Further emphasizing the relevance of these three markers in a neuroinflammatory setting, all three markers showed a significant correlation with neuroinflammatory marker CHI3L1 within the amyloid-positive AD group (Figure 6H).

## Discussion

Discrimination of microglial activity states is currently largely based on transcriptomic profiling.^11, 33, 35, 38, 41–44^ We previously reported opposite microglial phenotypes in mice lacking TREM2 or GRN. In the absence of TREM2 microglia were locked in a homeostatic state, whereas *Grn* ko microglia were hyperactivated.^11, 12^ We now made use of these findings and compared the opposite microglia signatures upon loss of TREM2 or GRN function on the proteome level in human and mouse models to explore potential markers for microglia activation and confirmed the translational applicability of our findings in CSF of AD and FTD-GRN patients.

The main finding of our study is the identification of six microglia activation-dependent markers, referred to as panel 6 including FABP3, MDH1, GDI1, CAPG, CD44, and GPNMB. These proteins are detectable and quantifiable in the conditioned media of hiMGL and in human CSF. Furthermore, they are all significantly increased in the conditioned media of *GRN* ko hiMGL, as well as in the CSF of FTD-GRN patients (Figure 3B). We decided not to refer to these changes as microglia-specific, as these proteins are expressed by other cell types as well. Nevertheless, due to the fact that the *GRN* ko hiMGL are grown in a monoculture with a confirmed activation signature and very little contamination from other brain cells, the significant upregulation of these proteins in the conditioned media of the *GRN* ko hiMGL strongly supports that the changes that we observed are indeed dependent on the microglial activation status. Furthermore, panel 6 proteins where also upregulated in FTD-GRN patients, where microglia are known to be hyperactivated.^13, 14^ Hence, we refer to our panel of six proteins as microglia activation-dependent. Our findings were further confirmed in a completely independent dataset derived from the EMIF-AD MBD cohort.^15^ CSF levels of FABP3, MDH1, and GDI1, were significantly elevated in MCI and AD patients of the EMIF-AD MDB cohort (Figure 6A-B). Strikingly, CSF levels of these three candidates were significantly different between MCI patients with and without detectable amyloid abnormalities (Figure 6D), thus the changes in their concentrations appear to be driven by early deposition of Aβ. This is in line with our previous findings in the DIAN cohort, where we found increased sTREM2 up to 21 years before the estimated year of symptom onset following immediately the deposition of amyloid.^7^ Together, these findings support the concept that microglia respond to the earliest amyloid related challenge even before profound cognitive changes are detected. Furthermore, our study revealed a significant correlation between CSF levels of FABP3, MDH1, and GDI1 in the FTD cohort (ALLFTD) and the AD cohort (EMIF-AD MBD) (Figure 5C and 6H). In the ALLFTD cohort, CSF levels of GPNMB showed a correlation with CSF TREM2, a correlation that is significant in FTD-GRN patients but not in healthy controls (Figure 5D-E). In addition, FABP3, MDH1, GDI1, CD44, and GPNMB individually correlated with the neuroinflammatory marker CHI3L1 (also known as YKL-40)^39, 40, 45, 46^, providing further evidence that the levels of these markers are indeed linked to a microglial response (i.e. a change in their activity state) (Figure 5C and 6H).

Further supporting our findings, abnormal CSF levels of individual members of panel 6 proteins have been reported in various neurodegenerative diseases: (1) FABP3 in cases of Parkinson’s disease with dementia (PDD), Dementia with Lewy bodies (DLB)^47^, AD^47–49^ and even in healthy individuals with confirmed amyloid pathology^50^, (2) MDH1 in sporadic (sCJD) and genetic Creutzfeldt-Jakob disease (gCJD)^51, 52^ and in AD^49^, (3) GDI1 in AD^49^, (4) CAPG in sporadic and genetic amyotrophic lateral sclerosis (ALS)^53^, (5) CD44 in AD^49^, and (6) GPNMB in sporadic and genetic amyotrophic lateral sclerosis (ALS)^53^.

Importantly, our data also revealed changes of the proteome that are not translatable across the mouse and human models, such as protein levels of MDH1, ENOPH1, CD84, and EPB41L3, proteins that are upregulated in the hiMGL model and in human CSF, but downregulated in mice (Appendix Figure S3). The translational commonalities and differences reported here may therefore also be of importance for selecting and understanding microglial responses in different models.

Biomarkers are not only relevant for clinical applications, but their changes may inform us about important metabolic abnormalities during disease onset and progression. FABP3 belongs to a family of proteins involved in uptake and metabolism of long-chain-fatty acids^54, 55^, while GPNMB appears to be involved in cell proliferation.^56^ Both may be highly relevant, since many if not all TREM2 agonistic antibodies boost microglial proliferation and ATV:TREM2, a brain penetrant anti-human TREM2 agonist also ameliorates lipid dysmetabolism.^3, 57^ Furthermore, levels of cellular FABP3 have been reported to increase in adipocytes and hepatocytes upon chronic exposure to high lipid concentrations.^54^ Finally, FABP3, together with APOE, were recently proposed as biomarkers for lipid metabolism in AD patients.^48^ It is therefore likely that the elevated FABP3 levels that we observe in microglia reflect a response to the lipid burden that microglia are facing in diseased conditions^58^ specifically upon GRN deficiency^59^ but also in other neurodegenerative diseases. MDH1 plays a central role in the malate-aspartate shuttle and the TCA cycle, which have been linked to LPS-induced microglial activation.^60^ Finally, GPNMB has been reported as highly expressed in microglial populations that either cluster around Aβ plaques or that are located distal to plaques but appear as amoeboid and with high lipid content.^61^ Of note, the soluble variant of GPNMB acts as a ligand for CD44, which is associated with anti-inflammatory responses as well as de novo lipogenesis.^62–64^ As previously suggested, lipid-related processes and metabolic reprogramming are of central importance in the context of microglial activation.^65–70^ Moreover, many of the protective functions of TREM2 have been linked to the sensing, binding, uptake, and metabolism of lipids, which are required for the microglial response.^71–75^ Of note, the currently used technology for in vivo assessing microglial activation makes use of translocator protein-positron emission tomography (TSPO-PET). TSPO-PET is based on the expression of TSPO protein, which is expressed at the mitochondrial membrane by multiple cell types throughout the body.^76^ TSPO is not microglia-specific, but appears to be microglia activation-dependent.^11, 77–80^ Interestingly, the main function of TSPO is to transport cholesterol across the mitochondrial membrane, where it can be further metabolized.^81^ Taken together, the microglial response is highly dependent on lipid-related processes, which further supports our idea that changes in lipid metabolism reflect microglia activation.

Due to the relative rarity of *GRN*-mutations associated with FTD (representing 5-20% of all FTD cases^82^), the rather small cohort size is a limitation of our study. In addition to the small cohort size, the obtained clinical data for the participants of the ALLFTD is very limited, which may affect the statistical analysis and compromise the interpretation of the ROC analysis. For these reasons, we employ this model as a companion to conventional tests to provide further evidence of the protein panel’s involvement, but not as exclusive evidence of the diagnostic capacities of this panel. Within the ALLFTD cohort and EMIF-AD MBD cohort Caucasian participants are overrepresented, which limits our study in the sense that none of these cohorts represent the global population. Therefore, additional human studies are required to confirm the selected candidates.

## Contributions

IP, CH, SAM and SFL designed the study. IP collected all samples from mouse and hiMGL. IP prepared all samples for label-free mass spectrometry analysis. SAM performed the mass spectrometry analysis. IP, SAM and AD performed the final statistical analysis. IP and CH interpreted the data. SR generated the hiPSC lines and performed the quality control and hiMGL differentiations under the supervision of DP. HZ and SFL supervised the mass spectrometry analysis. The original manuscript draft was predominantly written by IP and CH, with the help of SAM and AD. All authors reviewed and revised the manuscript.

## Declaration of Interests

CH collaborates with Denali Therapeutics. HZ has served at scientific advisory boards and/or as a consultant for Abbvie, Acumen, Alector, Alzinova, ALZPath, Annexon, Apellis, Artery Therapeutics, AZTherapies, CogRx, Denali, Eisai, Nervgen, Novo Nordisk, Optoceutics, Passage Bio, Pinteon Therapeutics, Prothena, Red Abbey Labs, reMYND, Roche, Samumed, Siemens Healthineers, Triplet Therapeutics, and Wave, has given lectures in symposia sponsored by Cellectricon, Fujirebio, Alzecure, Biogen, and Roche, and is a co-founder of Brain Biomarker Solutions in Gothenburg AB (BBS), which is a part of the GU Ventures Incubator Program (outside submitted work).

## Data sharing

The codes used for data sharing are available from IP, SAM, and SFL.

## Acknowledgements

This work was supported by the Deutsche Forschungsgemeinschaft (DFG, German Research Foundation) under Germany’s Excellence Strategy within the framework of the Munich Cluster for Systems Neurology (EXC 2145 SyNergy – ID 390857198 to CH, SFL and DP) and a Koselleck Project HA1737/16-1 (to CH). Further funding came from Alzheimer’s Association (to CH and DP), Vascular Dementia Research Foundation (to DP), and the donors of ADR AD2019604S, a program of the BrightFocus Foundation (to DP). HZ is a Wallenberg Scholar supported by grants from the Swedish Research Council (#2022-01018), the European Union’s Horizon Europe research and innovation programme under grant agreement No 101053962, Swedish State Support for Clinical Research (#ALFGBG-71320), the Alzheimer Drug Discovery Foundation (ADDF), USA (#201809-2016862), the AD Strategic Fund and the Alzheimer’s Association (#ADSF-21-831376-C, #ADSF-21-831381-C, and #ADSF-21-831377-C), the Bluefield Project, the Olav Thon Foundation, the Erling-Persson Family Foundation, Stiftelsen för Gamla Tjänarinnor, Hjärnfonden, Sweden (#FO2022-0270), the European Union’s Horizon 2020 research and innovation programme under the Marie Skłodowska-Curie grant agreement No 860197 (MIRIADE), the European Union Joint Programme – Neurodegenerative Disease Research (JPND2021-00694), and the UK Dementia Research Institute at UCL (UKDRI-1003).

We thank the staff and investigators of the ARTFL-LEFFTDS Longitudinal Frontotemporal Lobar Degeneration (ALLFTD) study (supported through a National Institute of Aging (NIA) and National Institute of Neurological Disorders and Stroke (NINDS) grant U19AG063911, as well as the participants and their families, whose help and participation made this work possible. The authors would like to thank Anna Berghofer for assisting with the preparation of samples for mass spectrometry analysis. Stephan Breimann for his help with the analysis of the MS-based data. Sophia Weiner, Johan Gobom, Maciej Dulewicz, and Juan Lantero Rodriguez for their help regarding the analysis of the data from the EMIF-AD MBD study. Samira Parhizkar for her expertise regarding CSF collection from mice and Laura Sebastian-Monasor for help with the isolation of microglia. We thank Michael Willem, Anne v. Thaden and Manuela Schneider for their help with mouse related work. We thank Oskar Hansson and Anja Capell for reviewing the manuscript.

## Appendix Tables and Figures

### Appendix Table S1: ALLFTD cohort demographics

Demographic data for non-carriers and mutation carriers.

### Appendix Table S2: Detailed description of the 26 proteins significantly increased in FTD-GRN CSF and media of *GRN* ko hiMGL

Description of the 26 proteins significantly upregulated in both CSF of FTD-GRN patients and in the conditioned media of *GRN* ko hiMGL. Information obtained from UniProt (uniprot.org on 21.07.2022).

### Appendix Figure S1: Generation of *TREM2* ko hiMGL

**(A)** CRISPR/Cas9 genome editing strategy of *TREM2* ko in hiPSCs: The *TREM2* locus was targeted in exon 2 by a sgRNA (target and PAM sequence indicated), leading to a one base pair insertion in the *TREM2*-knockout line. The resulting frameshift exposes a nearby stop codon. **(B)** mRNA levels of *TREM2* in WT and *TREM2*-knockout hiMGL, normalized to WT levels, measured by qPCR, showing that *TREM2* is not expressed on the RNA level in the *TREM2*-knockout (mean, SD). **(C)** sTREM2 concentration in conditioned media of WT and *TREM2*-knockout hiMGL, normalized to WT levels, measured by ELISA, showing no sTREM2 in the media of the *TREM2*-knockout. **(D)** Analysis of on-target effects mediated by CRISPR editing through Sanger sequencing of SNPs (as previously described^83^), near the edited locus in WT and *TREM2*-knockout lines, showing maintenance of both alleles. **(E)** Top five most similar off-target loci ranked by MIT and CFD prediction scores. No off-target effects were detected by Sanger sequencing. **(F)** Immunofluorescence image of the pluripotency markers SSEA4, NANOG, TRA160, and OCT4 with DAPI in *TREM2*-knockout iPSCs indicating no change in pluripotency. Scalebars = 100 µm. **(G)** No chromosomal aberrations were detected by molecular karyotyping: Log R ratios and B allele frequencies (BAF) for each chromosome in the *TREM2*-knockout iPSC line. The blue dots indicate measured SNPs. Normal zygosities on all chromosomes are indicated by BAF values and absence of detectable insertions and deletions are confirmed by the Log R ratios.

### Appendix Figure S2: Proteomic signature of hiMGL

**(A)** Comparing proteome and transcriptome changes of selected proteins in WT, *GRN* ko, and *TREM2* ko hiMGL. Confirming the activated signature in *GRN* ko hiMGL on a proteome level, a signature confirmed to be absent in the *TREM2* ko hiMGL. **(B-C)** Comparison of the proteomes of the *TREM2* ko hiMGL and *Trem2* ko mouse model. **(B)** Significantly upregulated proteins in cell lysates from *TREM2* ko hiMGL (pink) versus *Trem2* ko mice (orange). An overlap of 5 proteins (box: GLO1, PRPS1, HPRT1, SORT1, DHX36) were significantly upregulated in microglia of both *TREM2* ko systems. **(C)** Significantly upregulated proteins in conditioned media from *TREM2* ko hiMGL (pink) versus the CSF of *Trem2* ko mice (orange). The significantly upregulated changes in the secretome of *TREM2* ko hiMGL and the CSF from *Trem2* ko mice did not overlap. Significance was calculated using Student’s t-test, changes with p-value < 0·05 are considered significant.

### Appendix Figure S3: Binary comparisons of the 27 proteins significantly upregulated in *GRN* ko hiMGL secretome and FTD-GRN CSF

A binary comparison of the 26 proteins significantly upregulated in FTD-GRN CSF (hCSF) and in the secretome of *GRN* ko hiMGL (hiMGL M.) with the up-(red) or downregulation (blue) of each protein indicated for cell lysates of *GRN* ko hiMGL (hiMGL L.), CSF of *Grn* ko mice (mCSF), and in cell lysates of microglia isolated from *Grn* ko mice (mMG). Statistical differences were calculated using Student’s t-test. Crosses indicate missing values. * = p-value < 0·05, ** = p-value < 0·01, *** = p-value < 0·001, n.s. = non-significant. False discovery rate (FDR) was not considered for the presented visualizations. Mice used for analysis of the proteome were 12 months old. All presented proteomic data was obtained by LC-MS.

### Appendix Figure S4: The levels of panel 6 proteins in FTD-GRN CSF and conditioned media from *GRN* ko hiMGL

Abundances of panel 6 proteins: **(A)** FABP3, **(B)** MDH1, **(C)** GDI1, **(D)** CAPG, **(E)** CD44, and **(F)** GPNMB, in the media of *GRN*-knockout hiMGL (left graph) and the CSF of FTD-GRN patients (right graph), compared to control groups (WT or CON (= NC), respectively). *GRN* deficiency or mutation carriers are presented as yellow dots, with mild symptomatic FTD-GRN patients (mild cognitive impairment = MCI; n=3) presented as yellow triangles. Control groups are presented as gray dots. The mean of the respective control group is set to 100%. Statistical differences were calculated using Student’s t-test. * = p-value < 0·05, ** = p-value < 0·01, *** = p-value < 0·001, n.s. = non-significant. FDR was not considered for the presented visualizations.

### Appendix Figure S5: CSF levels of the panel 6 proteins in the EMIF-AD MBD cohort

CSF levels of CAPG (top), CD44 (middle) and GPNMB (bottom) in the EMIF-AD MBD cohort with **(A)** stratification according to diagnosis only and **(B)** stratification according to diagnosis and amyloid status. A^-^ = amyloid normal, A^+^ = amyloid abnormal.^15^ Significance was calculated using ordinary one-way ANOVA (alpha = 0·05).

## References

1. Romero-Molina C, Garretti F, Andrews SJ, Marcora E, Goate AM. Microglial efferocytosis: Diving into the Alzheimer’s disease gene pool. Neuron. 2022;110(21):3513–33.

2. Deczkowska A, Weiner A, Amit I. The Physiology, Pathology, and Potential Therapeutic Applications of the TREM2 Signaling Pathway. Cell. 2020;181(6):1207–17.

3. Lewcock JW, Schlepckow K, Di Paolo G, Tahirovic S, Monroe KM, Haass C. Emerging Microglia Biology Defines Novel Therapeutic Approaches for Alzheimer’s Disease. Neuron. 2020;108(5):801–21.

4. Ulland TK, Colonna M. TREM2 – a key player in microglial biology and Alzheimer disease. Nat Rev Neurol. 2018;14(11):667–75.

5. Kleinberger G, Yamanishi Y, Suarez-Calvet M, Czirr E, Lohmann E, Cuyvers E, et al. TREM2 mutations implicated in neurodegeneration impair cell surface transport and phagocytosis. Sci Transl Med. 2014;6(243):243ra86.

6. Suarez-Calvet M, Araque Caballero MA, Kleinberger G, Bateman RJ, Fagan AM, Morris JC, et al. Early changes in CSF sTREM2 in dominantly inherited Alzheimer’s disease occur after amyloid deposition and neuronal injury. Sci Transl Med. 2016;8(369):369ra178.

7. Morenas-Rodriguez E, Li Y, Nuscher B, Franzmeier N, Xiong C, Suarez-Calvet M, et al. Soluble TREM2 in CSF and its association with other biomarkers and cognition in autosomal-dominant Alzheimer’s disease: a longitudinal observational study. Lancet Neurol. 2022;21(4):329–41.

8. Parhizkar S, Arzberger T, Brendel M, Kleinberger G, Deussing M, Focke C, et al. Loss of TREM2 function increases amyloid seeding but reduces plaque-associated ApoE. Nat Neurosci. 2019;22(2):191–204.

9. Ewers M, Franzmeier N, Suarez-Calvet M, Morenas-Rodriguez E, Caballero MAA, Kleinberger G, et al. Increased soluble TREM2 in cerebrospinal fluid is associated with reduced cognitive and clinical decline in Alzheimer’s disease. Sci Transl Med. 2019;11(507).

10. Schlepckow K, Monroe KM, Kleinberger G, Cantuti-Castelvetri L, Parhizkar S, Xia D, et al. Enhancing protective microglial activities with a dual function TREM2 antibody to the stalk region. EMBO Mol Med. 2020;12(4):e11227.

11. Gotzl JK, Brendel M, Werner G, Parhizkar S, Sebastian Monasor L, Kleinberger G, et al. Opposite microglial activation stages upon loss of PGRN or TREM2 result in reduced cerebral glucose metabolism. EMBO Mol Med. 2019;11(6).

12. Mazaheri F, Snaidero N, Kleinberger G, Madore C, Daria A, Werner G, et al. TREM2 deficiency impairs chemotaxis and microglial responses to neuronal injury. EMBO Rep. 2017;18(7):1186–98.

13. Götzl JK, Mori K, Damme M, Fellerer K, Tahirovic S, Kleinberger G, et al. Common pathobiochemical hallmarks of progranulin-associated frontotemporal lobar degeneration and neuronal ceroid lipofuscinosis. Acta Neuropathol. 2014;127(6):845–60.

14. Lui H, Zhang J, Makinson SR, Cahill MK, Kelley KW, Huang HY, et al. Progranulin Deficiency Promotes Circuit-Specific Synaptic Pruning by Microglia via Complement Activation. Cell. 2016;165(4):921–35.

15. Weiner S, Sauer M, Visser PJ, Tijms BM, Vorontsov E, Blennow K, et al. Optimized sample preparation and data analysis for TMT proteomic analysis of cerebrospinal fluid applied to the identification of Alzheimer’s disease biomarkers. Clin Proteomics. 2022;19(1):13.

16. Kayasuga Y, Chiba S, Suzuki M, Kikusui T, Matsuwaki T, Yamanouchi K, et al. Alteration of behavioural phenotype in mice by targeted disruption of the progranulin gene. Behav Brain Res. 2007;185(2):110–8.

17. Turnbull IR, Gilfillan S, Cella M, Aoshi T, Miller M, Piccio L, et al. Cutting edge: TREM-2 attenuates macrophage activation. J Immunol. 2006;177(6):3520–4.

18. Lim NK, Moestrup V, Zhang X, Wang WA, Moller A, Huang FD. An Improved Method for Collection of Cerebrospinal Fluid from Anesthetized Mice. J Vis Exp. 2018(133).

19. Sebastian Monasor L, Muller SA, Colombo AV, Tanrioever G, Konig J, Roth S, et al. Fibrillar Abeta triggers microglial proteome alterations and dysfunction in Alzheimer mouse models. Elife. 2020;9.

20. Reifschneider A, Robinson S, van Lengerich B, Gnorich J, Logan T, Heindl S, et al. Loss of TREM2 rescues hyperactivation of microglia, but not lysosomal deficits and neurotoxicity in models of progranulin deficiency. EMBO J. 2022;41(4):e109108.

21. Tijms BM, Gobom J, Reus L, Jansen I, Hong S, Dobricic V, et al. Pathophysiological subtypes of Alzheimer’s disease based on cerebrospinal fluid proteomics. Brain. 2020;143(12):3776–92.

22. Wisniewski JR, Zougman A, Mann M. Combination of FASP and StageTip-based fractionation allows in-depth analysis of the hippocampal membrane proteome. J Proteome Res. 2009;8(12):5674–8.

23. Hughes CS, Sorensen PH, Morin GB. A Standardized and Reproducible Proteomics Protocol for Bottom-Up Quantitative Analysis of Protein Samples Using SP3 and Mass Spectrometry. Methods Mol Biol. 2019;1959:65–87.

24. Pigoni M, Wanngren J, Kuhn PH, Munro KM, Gunnersen JM, Takeshima H, et al. Seizure protein 6 and its homolog seizure 6-like protein are physiological substrates of BACE1 in neurons. Mol Neurodegener. 2016;11(1):67.

25. Rappsilber J, Ishihama Y, Mann M. Stop and go extraction tips for matrix-assisted laser desorption/ionization, nanoelectrospray, and LC/MS sample pretreatment in proteomics. Anal Chem. 2003;75(3):663–70.

26. Hughes CS, Moggridge S, Müller T, Sorensen PH, Morin GB, Krijgsveld J. Single-pot, solid-phase-enhanced sample preparation for proteomics experiments. Nat Protoc. 2019;14(1):68–85.

27. Tushaus J, Muller SA, Kataka ES, Zaucha J, Sebastian Monasor L, Su M, et al. An optimized quantitative proteomics method establishes the cell type-resolved mouse brain secretome. EMBO J. 2020;39(20):e105693.

28. Cox J, Hein MY, Luber CA, Paron I, Nagaraj N, Mann M. Accurate proteome-wide label-free quantification by delayed normalization and maximal peptide ratio extraction, termed MaxLFQ. Mol Cell Proteomics. 2014;13(9):2513–26.

29. Demichev V, Messner CB, Vernardis SI, Lilley KS, Ralser M. DIA-NN: neural networks and interference correction enable deep proteome coverage in high throughput. Nat Methods. 2020;17(1):41–4.

30. Tyanova S, Temu T, Sinitcyn P, Carlson A, Hein MY, Geiger T, et al. The Perseus computational platform for comprehensive analysis of (prote)omics data. Nat Methods. 2016;13(9):731–40.

31. Tusher VG, Tibshirani R, Chu G. Significance analysis of microarrays applied to the ionizing radiation response. Proc Natl Acad Sci U S A. 2001;98(9):5116–21.

32. Benjamini Y, Krieger AM, Yekutieli D. Adaptive linear step-up procedures that control the false discovery rate. Biometrika. 2006;93(3):491–507.

33. Deczkowska A, Keren-Shaul H, Weiner A, Colonna M, Schwartz M, Amit I. Disease-Associated Microglia: A Universal Immune Sensor of Neurodegeneration. Cell. 2018;173(5):1073–81.

34. Holtman IR, Raj DD, Miller JA, Schaafsma W, Yin Z, Brouwer N, et al. Induction of a common microglia gene expression signature by aging and neurodegenerative conditions: a co-expression meta-analysis. Acta Neuropathol Commun. 2015;3:31.

35. Keren-Shaul H, Spinrad A, Weiner A, Matcovitch-Natan O, Dvir-Szternfeld R, Ulland TK, et al. A Unique Microglia Type Associated with Restricting Development of Alzheimer’s Disease. Cell. 2017;169(7):1276–90 e17.

36. Krasemann S, Madore C, Cialic R, Baufeld C, Calcagno N, El Fatimy R, et al. The TREM2-APOE Pathway Drives the Transcriptional Phenotype of Dysfunctional Microglia in Neurodegenerative Diseases. Immunity. 2017;47(3):566–81 e9.

37. Holtman IR, Skola D, Glass CK. Transcriptional control of microglia phenotypes in health and disease. J Clin Invest. 2017;127(9):3220–9.

38. Butovsky O, Weiner HL. Microglial signatures and their role in health and disease. Nat Rev Neurosci. 2018;19(10):622–35.

39. Janelidze S, Hertze J, Zetterberg H, Landqvist Waldo M, Santillo A, Blennow K, et al. Cerebrospinal fluid neurogranin and YKL-40 as biomarkers of Alzheimer’s disease. Ann Clin Transl Neurol. 2016;3(1):12–20.

40. Woollacott IOC, Nicholas JM, Heller C, Foiani MS, Moore KM, Russell LL, et al. Cerebrospinal Fluid YKL-40 and Chitotriosidase Levels in Frontotemporal Dementia Vary by Clinical, Genetic and Pathological Subtype. Dement Geriatr Cogn Disord. 2020;49(1):56–76.

41. Grubman A, Choo XY, Chew G, Ouyang JF, Sun G, Croft NP, et al. Transcriptional signature in microglia associated with Abeta plaque phagocytosis. Nat Commun. 2021;12(1):3015.

42. Hammond TR, Dufort C, Dissing-Olesen L, Giera S, Young A, Wysoker A, et al. Single-Cell RNA Sequencing of Microglia throughout the Mouse Lifespan and in the Injured Brain Reveals Complex Cell-State Changes. Immunity. 2019;50(1):253–71 e6.

43. Matcovitch-Natan O, Winter DR, Giladi A, Vargas Aguilar S, Spinrad A, Sarrazin S, et al. Microglia development follows a stepwise program to regulate brain homeostasis. Science. 2016;353(6301):aad8670.

44. Sankowski R, Bottcher C, Masuda T, Geirsdottir L, Sagar, Sindram E, et al. Mapping microglia states in the human brain through the integration of high-dimensional techniques. Nat Neurosci. 2019;22(12):2098–110.

45. Llorens F, Thune K, Tahir W, Kanata E, Diaz-Lucena D, Xanthopoulos K, et al. YKL-40 in the brain and cerebrospinal fluid of neurodegenerative dementias. Mol Neurodegener. 2017;12(1):83.

46. Craig-Schapiro R, Perrin RJ, Roe CM, Xiong C, Carter D, Cairns NJ, et al. YKL-40: a novel prognostic fluid biomarker for preclinical Alzheimer’s disease. Biol Psychiatry. 2010;68(10):903–12.

47. Chiasserini D, Biscetti L, Eusebi P, Salvadori N, Frattini G, Simoni S, et al. Differential role of CSF fatty acid binding protein 3, alpha-synuclein, and Alzheimer’s disease core biomarkers in Lewy body disorders and Alzheimer’s dementia. Alzheimers Res Ther. 2017;9(1):52.

48. Dulewicz M, Kulczynska-Przybik A, Slowik A, Borawska R, Mroczko B. Fatty Acid Binding Protein 3 (FABP3) and Apolipoprotein E4 (ApoE4) as Lipid Metabolism-Related Biomarkers of Alzheimer’s Disease. J Clin Med. 2021;10(14).

49. Higginbotham L, Ping L, Dammer EB, Duong DM, Zhou M, Gearing M, et al. Integrated proteomics reveals brain-based cerebrospinal fluid biomarkers in asymptomatic and symptomatic Alzheimer’s disease. Sci Adv. 2020;6(43).

50. Dhiman K, Villemagne VL, Fowler C, Bourgeat P, Li QX, Collins S, et al. Cerebrospinal fluid levels of fatty acid-binding protein 3 are associated with likelihood of amyloidopathy in cognitively healthy individuals. Alzheimers Dement (Amst). 2022;14(1):e12377.

51. Schmitz M, Llorens F, Pracht A, Thom T, Correia A, Zafar S, et al. Regulation of human cerebrospinal fluid malate dehydrogenase 1 in sporadic Creutzfeldt-Jakob disease patients. Aging (Albany NY). 2016;8(11):2927–35.

52. Zerr I, Villar-Pique A, Schmitz VE, Poleggi A, Pocchiari M, Sanchez-Valle R, et al. Evaluation of Human Cerebrospinal Fluid Malate Dehydrogenase 1 as a Marker in Genetic Prion Disease Patients. Biomolecules. 2019;9(12).

53. Oeckl P, Weydt P, Thal DR, Weishaupt JH, Ludolph AC, Otto M. Proteomics in cerebrospinal fluid and spinal cord suggests UCHL1, MAP2 and GPNMB as biomarkers and underpins importance of transcriptional pathways in amyotrophic lateral sclerosis. Acta Neuropathol. 2020;139(1):119–34.

54. Furuhashi M, Hotamisligil GS. Fatty acid-binding proteins: role in metabolic diseases and potential as drug targets. Nat Rev Drug Discov. 2008;7(6):489–503.

55. Binas B, Danneberg H, McWhir J, Mullins L, Clark AJ. Requirement for the heart-type fatty acid binding protein in cardiac fatty acid utilization. FASEB J. 1999;13(8):805–12.

56. Li Q, Cheng Z, Zhou L, Darmanis S, Neff NF, Okamoto J, et al. Developmental Heterogeneity of Microglia and Brain Myeloid Cells Revealed by Deep Single-Cell RNA Sequencing. Neuron. 2019;101(2):207–23.e10.

57. van Lengerich B, Zhan L, Xia D, Chan D, Joy D, Park JI, et al. A TREM2-activating antibody with a blood–brain barrier transport vehicle enhances microglial metabolism in Alzheimer’s disease models. Nature Neuroscience. 2023;26(3):416–29.

58. Munoz Herrera OM, Zivkovic AM. Microglia and Cholesterol Handling: Implications for Alzheimer’s Disease. Biomedicines. 2022;10(12).

59. Logan T, Simon MJ, Rana A, Cherf GM, Srivastava A, Davis SS, et al. Rescue of a lysosomal storage disorder caused by Grn loss of function with a brain penetrant progranulin biologic. Cell. 2021;184(18):4651–68.e25.

60. Zhou C, Shang W, Yin S-K, Shi H, Ying W. Malate-Aspartate Shuttle Plays an Important Role in LPS-Induced Neuroinflammation of Mice Due to its Effect on STAT3 Phosphorylation. Frontiers in Molecular Biosciences. 2021;8.

61. Huttenrauch M, Ogorek I, Klafki H, Otto M, Stadelmann C, Weggen S, et al. Glycoprotein NMB: a novel Alzheimer’s disease associated marker expressed in a subset of activated microglia. Acta Neuropathol Commun. 2018;6(1):108.

62. Kuang H, Lin JD. GPNMB: expanding the code for liver-fat communication. Nat Metab. 2019;1(5):507–8.

63. Neal ML, Boyle AM, Budge KM, Safadi FF, Richardson JR. The glycoprotein GPNMB attenuates astrocyte inflammatory responses through the CD44 receptor. J Neuroinflammation. 2018;15(1):73.

64. Saade M, Araujo de Souza G, Scavone C, Kinoshita PF. The Role of GPNMB in Inflammation. Front Immunol. 2021;12:674739.

65. Chausse B, Kakimoto PA, Kann O. Microglia and lipids: how metabolism controls brain innate immunity. Semin Cell Dev Biol. 2021;112:137–44.

66. Loving BA, Bruce KD. Lipid and Lipoprotein Metabolism in Microglia. Front Physiol. 2020;11:393.

67. Paolicelli RC, Sierra A, Stevens B, Tremblay ME, Aguzzi A, Ajami B, et al. Microglia states and nomenclature: A field at its crossroads. Neuron. 2022;110(21):3458–83.

68. Kunkle BW, Grenier-Boley B, Sims R, Bis JC, Damotte V, Naj AC, et al. Genetic meta-analysis of diagnosed Alzheimer’s disease identifies new risk loci and implicates Abeta, tau, immunity and lipid processing. Nat Genet. 2019;51(3):414–30.

69. Leyrolle Q, Laye S, Nadjar A. Direct and indirect effects of lipids on microglia function. Neurosci Lett. 2019;708:134348.

70. Churchward MA, Tchir DR, Todd KG. Microglial Function during Glucose Deprivation: Inflammatory and Neuropsychiatric Implications. Mol Neurobiol. 2018;55(2):1477–87.

71. Ulland TK, Song WM, Huang SC, Ulrich JD, Sergushichev A, Beatty WL, et al. TREM2 Maintains Microglial Metabolic Fitness in Alzheimer’s Disease. Cell. 2017;170(4):649–63 e13.

72. Wang Y, Cella M, Mallinson K, Ulrich JD, Young KL, Robinette ML, et al. TREM2 lipid sensing sustains the microglial response in an Alzheimer’s disease model. Cell. 2015;160(6):1061–71.

73. Poliani PL, Wang Y, Fontana E, Robinette ML, Yamanishi Y, Gilfillan S, et al. TREM2 sustains microglial expansion during aging and response to demyelination. J Clin Invest. 2015;125(5):2161–70.

74. Nugent AA, Lin K, van Lengerich B, Lianoglou S, Przybyla L, Davis SS, et al. TREM2 Regulates Microglial Cholesterol Metabolism upon Chronic Phagocytic Challenge. Neuron. 2020;105(5):837–54 e9.

75. Gouna G, Klose C, Bosch-Queralt M, Liu L, Gokce O, Schifferer M, et al. TREM2-dependent lipid droplet biogenesis in phagocytes is required for remyelination. J Exp Med. 2021;218(10).

76. Kreisl WC, Henter ID, Innis RB. Imaging Translocator Protein as a Biomarker of Neuroinflammation in Dementia. Adv Pharmacol. 2018;82:163–85.

77. Beckers L, Ory D, Geric I, Declercq L, Koole M, Kassiou M, et al. Increased Expression of Translocator Protein (TSPO) Marks Pro-inflammatory Microglia but Does Not Predict Neurodegeneration. Mol Imaging Biol. 2018;20(1):94–102.

78. Kleinberger G, Brendel M, Mracsko E, Wefers B, Groeneweg L, Xiang X, et al. The FTD-like syndrome causing TREM2 T66M mutation impairs microglia function, brain perfusion, and glucose metabolism. EMBO J. 2017;36(13):1837–53.

79. Pozzo ED, Tremolanti C, Costa B, Giacomelli C, Milenkovic VM, Bader S, et al. Microglial Pro-Inflammatory and Anti-Inflammatory Phenotypes Are Modulated by Translocator Protein Activation. Int J Mol Sci. 2019;20(18).

80. Xiang X, Wind K, Wiedemann T, Blume T, Shi Y, Briel N, et al. Microglial activation states drive glucose uptake and FDG-PET alterations in neurodegenerative diseases. Sci Transl Med. 2021;13(615):eabe5640.

81. Papadopoulos V, Miller WL. Role of mitochondria in steroidogenesis. Best Pract Res Clin Endocrinol Metab. 2012;26(6):771–90.

82. Amin S, Carling G, Gan L. New insights and therapeutic opportunities for progranulin-deficient frontotemporal dementia. Current Opinion in Neurobiology. 2022;72:131–9.

83. Weisheit I, Kroeger JA, Malik R, Klimmt J, Crusius D, Dannert A, et al. Detection of Deleterious On-Target Effects after HDR-Mediated CRISPR Editing. Cell Rep. 2020;31(8):107689.

